# Structural and functional characterization of the Severe fever with thrombocytopenia syndrome virus L protein

**DOI:** 10.1101/2020.03.02.973065

**Authors:** Dominik Vogel, Sigurdur Rafn Thorkelsson, Emmanuelle R. J. Quemin, Kristina Meier, Tomas Kouba, Nadja Gogrefe, Carola Busch, Sophia Reindl, Stephan Günther, Stephen Cusack, Kay Grünewald, Maria Rosenthal

**Author notes:** To whom correspondence should be addressed: Maria Rosenthal. These authors contributed equally to this work (Dominik Vogel, Sigurdur Rafn Thorkelsson).

## Abstract

The *Bunyavirales* order contains several emerging viruses with high epidemic potential, including Severe fever with thrombocytopenia syndrome virus (SFTSV). The lack of medical countermeasures, such as vaccines and antivirals, is a limiting factor for the containment of any virus outbreak. To develop such antivirals a profound understanding of the viral replication process is essential. The L protein of bunyaviruses is a multi-functional and multi-domain protein performing both virus transcription and genome replication and, therefore, would be an ideal drug target. We established expression and purification procedures for the full-length L protein of SFTSV. By combining single-particle electron-cryo microscopy and X-ray crystallography, we obtained 3D models covering ∼70% of the SFTSV L protein in the apo-conformation including the polymerase core region, the endonuclease and the cap-binding domain. We compared this first L structure of the *Phenuiviridae* family to the structures of La Crosse peribunyavirus L protein and influenza orthomyxovirus polymerase. Together with a comprehensive biochemical characterization of the distinct functions of SFTSV L protein, this work provides a solid framework for future structural and functional studies of L protein-RNA interactions and the development of antiviral strategies against this group of emerging human pathogens.

## INTRODUCTION

Severe fever with thrombocytopenia syndrome virus (SFTSV) is prevalent in East Asia and closely related to the Heartland virus that has been isolated in the US (1). Very recently an SFTSV-like virus has also been detected in bats from Germany (2). SFTSV belongs to the *Bunyavirales* order, established in 2018 to accommodate formerly separated bunya- and arenaviruses (3). Bunyaviruses are a diverse group of viruses with a segmented single-stranded RNA genome of negative polarity. They are distributed worldwide causing zoonoses with several outbreaks annually, mainly occurring in Low-to-Middle-Income countries and primarily affecting poor populations with restricted access to health care. Therefore, several bunyaviruses are listed on the WHO R&D Blueprint (4), a global strategy to enhance preparedness to future epidemics urging for the development of medical countermeasures, such as vaccines and antivirals, which are largely lacking. To develop antiviral strategies against emerging viruses, including SFTSV and other bunyaviruses a profound understanding of the viral life cycle is essential, especially of the similarities and differences within this virus group. The key player in bunyavirus transcription and genome replication is the large (L) protein with a size between 250 and 450 kDa, which contains multiple domains and functions including the viral RNA-dependent RNA polymerase (RdRp). For influenza virus, another segmented negative sense RNA virus (sNSV), small molecules targeting different functions of the polymerase complex have been clinically approved (5,6). However, in contrast to influenza viruses, whose polymerase has been extensively investigated, structural and functional information about bunyavirus L proteins are scarce.

During the processes of genome replication and transcription, bunyavirus L protein synthesizes three distinct RNA species: (i) antigenomic <u>c</u>omplementary RNA (cRNA), (ii) genomic <u>v</u>iral RNA (vRNA), and (iii) capped, mostly non-polyadenylated viral mRNA. Whereas genome replication is believed to be initiated *de novo* by the L protein, viral transcription is dependent on short, capped RNA primers derived from cellular RNAs by a mechanism called cap-snatching (7), which involves cap binding and endonuclease functions. The part of bunyavirus L protein that has been most characterized, both structurally and functionally, is the ∼20 kDa endonuclease domain (7-9). In most bunyavirus L proteins, this domain is located at the N terminus. Its metal-dependent RNA cleavage activity is needed for the cap-snatching mechanism to steal 5’ cap structures from cellular RNA and attach these fragments to the viral mRNA in order to be processed by the cellular translation machinery at the ribosome. For Rift Valley Fever virus (RVFV), a bunyavirus of the *Phenuiviridae* family, the C-terminal region of the L protein has been shown to contain a cap-binding domain (CBD), the second activity needed for the cap-snatching mechanism. This CBD is structurally similar to that of influenza virus but with distinct differences in its mode of cap-recognition (10,11). The third important activity is the RdRp, which is needed for both transcription and genome replication. To date, only for one bunyavirus L protein, that of La Crosse virus (LACV, *Peribunyaviridae* family), more extensive high-resolution structural information is available. The LACV structures contain ∼77% of the L protein sequence including the RdRp domain and important protein–RNA interaction sites but are entirely missing the C-terminal domain (12). Even though further structural insights into distinct functional states of bunyavirus L protein are still lacking, this was a milestone on the path to a complete structural understanding of bunyavirus transcription and replication. In terms of functional data, biochemical characterization of Lassa and Machupo viruses (*Arenaviridae* family) (13-16) as well as LACV (12) L proteins have been published. Specific to each bunyavirus family are the 3’ and 5’ termini of the genomic RNA segments that are highly conserved and almost fully complementary in sequence, denoted as the conserved promoter regions. For influenza virus and LACV, the RNA 5’ end forms a hook-like secondary structure that is bound in a specific pocket adjacent to the RNA synthesis active site of the L protein (12,17,18). The presence of a hook-conformation and its importance for robust RdRp activity have also been proposed for arenaviruses (13,16). Furthermore, for arenaviruses, it was shown that the L protein initiates genome synthesis *de novo* by a prime-and-realign mechanism (13) similar to the initiation of influenza virus vRNA synthesis and Hantaan virus genome replication (19-22). In order to deepen our understanding of bunyavirus genome replication and transcription, we aimed to structurally and functionally characterize the full-length L protein of a virus of the *Phenuiviridae* family. For this virus family, no high-resolution structural data on the L protein core domain is currently available. We established expression and purification of the full-length SFTSV L protein in insect cells using a baculovirus expression system with a yield of up to 11 mg per liter of culture. We present structural data on the RdRp core and adjacent L protein domains of SFTSV bunyavirus (*Phenuiviridae* family) obtained by electron cryo-microscopy (cryo-EM) and X-ray crystallography. Although we were not able to build the N-terminal endonuclease domain *de novo*, we could fit the recently published SFTSV endonuclease crystal structure (23) into the cryo-EM density map we obtained, thereby proposing an integrated model, which was also verified using data from small-angle X-ray scattering (SAXS) experiments. While the C-terminal region of the L protein was not resolved in the cryo-EM map, we solved the structure of the SFTSV CBD in complex with a cap-analogue by X-ray crystallography with a resolution of 1.35 Å and characterized how it interacts with cap-structures using biochemical and biophysical assays. We further used biochemical assays to characterize the different L protein activities, such as promoter binding, endonuclease and polymerase activities. In summary, we provide novel structural and functional information on SFTSV L protein that can serve as a basis for an improved understanding of bunyavirus transcription and genome replication processes and aid drug development.

## MATERIALS AND METHODS

### Cloning, expression and purification of SFTSV full-length L protein

The L gene of SFTSV strain AH12 (accession no. HQ116417) with a C-terminal StrepII-tag was chemically synthesized (Centic Biotech, Germany) and the plasmid-integrated gene was amplified via PCR. In the same step, primers were used for the introduction of mutations to the gene in the case of the catalytically inactive L proteins (D112A and D1126A). Amplified genes were cloned into an altered pFastBacHT B vector using the In-Fusion HD EcoDry Cloning Kit (Clontech). After transformation of DH10EMBacY *E. coli* cells (kindly provided by Imre Berger), which contain a bacmid as well as a plasmid coding for a topoisomerase, with the pFastBac-plasmids, recombinant bacmids were isolated and transfected into Sf21 insect cells for recombinant baculovirus production. Hi5 insect cells were used for the expression of the StrepII-tagged L proteins. The harvested cells were resuspended in buffer A (50 mM HEPES(NaOH) pH 7.0, 1 M NaCl, 10% (w/w) Glycerol and 2 mM dithiothreitol), supplemented with 0.05% (v/v) Tween20 and protease inhibitors (Roche, cOmplete mini), lysed by sonication and centrifuged two times at 20,000 x g for 30 min at 4°C. Soluble protein was loaded on Strep-TactinXT beads (IBA) and eluted with 50 mM Biotin (Applichem) in buffer B (50 mM HEPES(NaOH) pH 7.0, 500 mM NaCl, 10% (w/w) Glycerol and 2 mM dithiothreitol). L protein-containing fractions were pooled and diluted with an equal volume of buffer C (20 mM HEPES(NaOH) pH 7.0) before loading onto a heparin column (HiTrap Heparin HP, GE Healthcare). Proteins were eluted with buffer A and concentrated using centrifugal filter units (Amicon Ultra, 30 kDa MWCO). The proteins were further purified by size-exclusion chromatography (SD200, GE Healthcare) in buffer B and either used for biochemical assays or SAXS experiments. Purified L proteins were concentrated as described above, flash frozen in liquid nitrogen and stored at -80°C.

### Cloning, expression and purification of SFTSV cap-binding domain

The L gene region corresponding to amino acid residues 1695 to 1810 of SFTSV strain AH12 (accession no. HQ116417) was cloned into a pOPINF vector (24) using the NEBuilder HiFi DNA Assembly Cloning Kit (New England BioLabs). The protein, which based on sequence alignments corresponds to the CBD of RVFV (11) and therefore referred to as SFTSV CBD, was expressed as a wild-type version or mutated version (with single amino acid exchanges F1705A, Q1707A or Y1719A) in *E. coli* strain BL21 Gold (DE3) (Novagen) at 17°C overnight using TB medium and 0.5 mM isopropyl-β-D-thiogalactopyranosid for induction. After pelleting, the cells were resuspended in 50 mM Na-phosphate pH 7.5, 100 mM NaCl, 10 mM imidazole, Complete protease inhibitor EDTA-free (Roche), 0.4% (v/v) Triton X-100 and 0.025% (w/v) lysozyme and subsequently disrupted by sonication. The protein was purified from the soluble fraction after centrifugation by Ni affinity chromatography. Washing buffers contained 50 mM imidazole and 1 M NaCl or 50 mM imidazole and 100 mM NaCl. Elution buffer contained 500 mM imidazole and the eluted protein was immediately diluted with 20 mM Na-phosphate pH 6.5 followed by passing through an anion exchange chromatography column (HiTrap Q FF, GE Healthcare) and subsequent cation exchange chromatography (HiTrap SP FF, GE Healthcare, loading buffer: 50 mM Na-phosphate pH 6.5, 50 mM NaCl, 10% (w/v) glycerol, elution with salt gradient up to 1 M NaCl). The next step was the removal of the N-terminal His-tag by a GST-tagged 3C protease at 4°C overnight. After addition of up to 3 mM m^7^GTP the protein was concentrated for a final size exclusion chromatography (Superdex 200, 50 mM Na-phosphate, pH 6.5, 150 mM NaCl, 10% (w/v) glycerol). Purified proteins were concentrated using centrifugal devices with addition of up to 4 mM m^7^GTP (final concentration), flash frozen in liquid nitrogen, and stored at -80°C. For thermal shift assays and isothermal titration calorimetry, the purification procedure was done as described above but without any addition of m^7^GTP.

### Isothermal titration calorimetry

Affinity of SFTSV CBD to m^7^GTP and GTP was measured by isothermal titration calorimetry (ITC) using a MicroCal PEAQ-ITC instrument (Malvern Panalytical). Proteins were dialyzed overnight at 4°C against 50 mM Na-phosphate pH 6.5, 150 mM NaCl, 10% (w/v) glycerol. Ligand m^7^GTP was dissolved and GTP was diluted in the exact same dialysis buffer. Titrations were done with 150 µM CBD in the cell and 5.0-6.5 mM m^7^GTP or GTP in the syringe at 25°C with 19 injections of 2 µl (first injection 0.5 µl). Spacing between injections was constant with 150 seconds for all measurements. Data were analyzed and fitted with the respective PEAQ ITC evaluation software (Malvern) applying a single side binding model and fixing the stoichiometry value to 1.

### Thermal stability assay

Thermal stability of SFTSV CBD was measured by thermofluor assay (25). The assay contained a final concentration of 8 µM of CBD protein, 20 mM Na-phosphate pH 6.5, 100 mM NaCl, SYPRO-Orange (final dilution 1:1000) and either no additive or between 1 and 10 mM of m^7^GTP, m^7^GpppG, GTP or ATP. Thermal stability of influenza A virus PB2 CBD and RVFV CBD was assessed at a final protein concentration of 10 µM and 8 µM, respectively, in the same setup as described for SFTSV CBD.

### Electrophoretic mobility shift assay

RNAs (Supplementary Table S1) were chemically synthesized (Biomers) and labelled with T4 polynucleotide kinase (Thermo Fisher) and γ^32^P-ATP (Hartman Analytic), or ScriptCap m^7^G Capping System (CellScript) and α^32^P-GTP. Labeled RNA substrates were subsequently separated from unincorporated γ^32^P using Microspin G25 columns (GE Healthcare) or ethanol precipitation. RNA was heated for 3 min at 95°C and cooled down on ice to give single-stranded RNA (ssRNA). Reactions containing 0-14 pmol L protein and 2 pmol ^32^P-labelled ssRNA were set up in 10 μL binding buffer (50 mM HEPES(NaOH) pH 7.0, 150 mM NaCl, 5 mM MgCl_2_, 2 mM dithiothreitol, 10% glycerol, 0.5 μg/μL Poly(C) RNA (Sigma), 0.5 μg/μL bovine serum albumin and 0.5 U/μl RNasin (Promega)). After incubation on ice for 30 min, the products were separated by native gel electrophoresis using 6% polyacrylamide Tris-glycine gels and Tris-glycine buffer on ice. Signals were visualized by phosphor screen autoradiography using a Typhoon scanner (GE Healthcare) and quantified using ImageJ software (61) if necessary.

### Endonuclease Assay

Poly-A RNA 27mer and 40mer were chemically synthesized (Biomers) and ^32^P-labelled with T4 polynucleotide kinase (Thermo Fisher) or ScriptCap m7G Capping System (CellScript). RNA substrates were subsequently purified with a Microspin G25 column (GE Healthcare). Reactions containing 2.5 pmol protein and 3 pmol ^32^P-labelled RNA were carried out in a volume of 10 µL with 0.5 U/μl RNasin (Promega), 100 mM HEPES(NaOH) pH 7.0, 100 mM NaCl, 50 mM KCl, 2 mM MnCl_2_, 1 mM dithiothreitol, and 0.1 µg/µL bovine serum albumin and incubated at 37°C for 30 min. The reaction was stopped by adding an equivalent volume of RNA loading buffer (98% formamide, 18 mM EDTA, 0.025 mM SDS, xylene cyanol and bromophenol blue) and heating the samples at 95°C for 3 min. Products were separated by electrophoresis on denaturing 7 M urea, 20% polyacrylamide Tris-borate-EDTA gels in 0.5-fold Tris-borate buffer. Signals were visualized by phosphor screen autoradiography using a Typhoon scanner (GE Healthcare).

### Polymerase Assay

If not indicated otherwise, 5 pmol L protein were preincubated for 15 min with 6 pmol of 5’ promoter ssRNA (Supplementary Table S1) in assay buffer (100 mM HEPES(NaOH) pH 7.0, 100 mM NaCl, 50 mM KCl, 5 mM MgCl_2_, 0.5 U/μl RNasin (Promega), 2 mM dithiothreitol, and 0.1 μg/μL bovine serum albumin). Subsequently, 6 pmol of 3’ promoter ssRNA (Supplementary Table S1) and NTPs (0.8 mM UTP/ATP/CTP and 0.5 mM GTP supplemented with 5 μCi [α]^32^P-GTP) were added to a final reaction volume of 10 μL. After 1h at 30°C the reaction was stopped by adding an equivalent volume of RNA loading buffer (98% formamide, 18 mM EDTA, 0.025 mM SDS, xylene cyanol and bromophenol blue) and heating the sample for 5 min at 98°C. Products were separated by electrophoresis on denaturing 7 M urea, 20% polyacrylamide Tris-borate-EDTA gels in 0.5-fold Tris-borate buffer. Signals were visualized by phosphor screen autoradiography using a Typhoon scanner (GE Healthcare).

### Double-stranded RNA separation assay

RNA oligonucleotides (Supplementary Table S1) were chemically synthesized (Biomers) and 3’ labeled with T4 RNA ligase (Thermo Fisher) and pCp-Cy3 (Jena Bioscience). Labeled RNA substrates were subsequently separated from excess pCp-Cy3 by ethanol precipitation. RNA was mixed in 1:1 ratio in 100 mM HEPES(NaOH) pH 7.0, 100 mM NaCl, 50 mM KCl, 5 mM MgCl_2_, 0.5 U/μl RNasin (Promega), 2 mM dithiothreitol, and 0.1 μg/μL bovine serum albumin and incubated at 30°C for 30 min before an equal volume of RNA loading buffer (98% formamide, 18 mM EDTA, 0.025 mM SDS) was added. The samples were heated at 95°C for 3 min before separation on denaturing 7M Urea, 20% polyacrylamide Tris-borate-EDTA gels and 0.5-fold Tris-borate buffer.

### Crystallisation and crystallography

SFTSV CBD crystals grew within 14 days after mixing of one part SFTSV CBD at 18.6 mg/mL protein concentration supplemented with 4 mM m^7^GTP in 50 mM Na-phosphate, pH 6.5, 150 mM NaCl, 10% (w/v) glycerol with two parts of 7% (v/v) 2-Butanol, 150 mM MES pH 6.0 and 27.4% PEG 4000 in a sitting drop vapor diffusion setup at 20°C. Crystals were flash frozen in liquid nitrogen without cryo protectants and datasets were obtained at beamline P14 of PETRA III at Deutsches Elektronen Synchrotron (DESY), Hamburg, Germany. Datasets were processed with XDS (26) and the SFTSV CBD structure was solved by molecular replacement using Phaser (27) and the RVFV CBD (PDB 6QHG) monomer without the β-hairpin and loops as an input model. The structure was refined by iterative cycles of manual model building in Coot (28) and computational optimization with PHENIX (29). Visualization of structural data was done using either the PyMOL Molecular Graphics System, Version 1.7 Schrödinger, LLC or UCSF Chimera (30).

### Electron-cryo microscopy

Aliquots of 3 µL of SFTSV L protein diluted to 0.6 μM in buffer (20 mM HEPES pH 7.0, 500 mM NaCl, 20 mM MgCl_2_) were applied to glow-discharged Quantifoil R 2/1 Au G200F4 grids, immediately blotted for 2 s using an FEI Vitrobot Mk IV (4°C, 100% humidity, blotting force -10) and plunge frozen in liquid ethane/propane cooled to liquid nitrogen temperature. The grids were loaded into a 300-keV Titan Krios transmission electron cryo-microscope (Thermo Scientific) equipped with a K3 direct electron detector and a GIF BioQuantum energy filter (Gatan). A total of 2,626 movie frames were collected using the EPU software (Thermo Scientific) at a nominal magnification of x105 000 with a pixel size of 0.87 Å and a defocus range of -1 to -3 μm. Each movie fractionated into 60 frames, was collected for a total exposure time of 3 s with a flux of 15 electrons per physical pixel per second and a total dose of 59.45 e/Å^2^ with a 20-eV slit for the GIF (Supplementary Table S2).

### Cryo-EM data processing and model building

All movie frames were aligned using MotionCor2 and particles were picked automatically using Warp (31). The motion-corrected micrographs were imported in Relion 3.0 (32) and used for contrast transfer function (CTF) parameter calculation with gctf (33). The particles were extracted using coordinates from Warp (∼780,000) and subjected to two iterative rounds of reference-free 2D classification. Selected 2D classes representing the different projection views (575,157 particles) were initially classified into six 3D classes with image alignment using an *ab initio* volume created in cisTEM (34) and low-pass filtered to 40 Å, as a reference. The 3D classification with regularization parameter T = 4 was performed with 7.5° angular sampling for the first 25 iterations followed by 10 additional iterations using 3.7° angular sampling. One major class was identified as the most populated (223,604 particles) and better defined class, and was selected for auto-refinement and post-processing in Relion resulting in a 3D electron density map (SFTSV map A) with an estimated resolution of 3.8 Å using gold standard Fourier Shell Correlation (0.143 cut-off). The globally refined volume was then used as a template to sort particles again into 10 new 3D classes. This time a distinct class was visible containing additional volume for the C-terminal region (73,357 particles). This class submitted to auto-refine and post-processing and yielded a 3D EM map of 4.3 Å estimated resolution. The model was built *de novo* into SFTSV map A by the Map-to-model program included in Phenix (35) using the core domain of LACV L protein structure (PDB 5AMQ) as a starting model. The structure was refined by iterative cycles of manual model building in Coot (28) and computational optimization with PHENIX (29).

### Small angle X-ray scattering

Small angle X-ray scattering (SAXS) of SFTSV L protein was performed with an in-line size exclusion chromatography on a Superdex 200 Increase 5/150 GL column with a buffer containing 50 mM HEPES(NaOH) pH 7, 500 mM NaCl, 5% (w/v) glycerol, and 2 mM dithiothreitol. Data was collected at the SAXS beamline P12 of PETRA III storage ring of the DESY, Hamburg, Germany (36) using a PILATUS 2M pixel detector at 3.1 m sample distance and 10 keV energy (λ = 1.24 Å), a momentum transfer range of 0.01 Å^-1^ < s < 0.45 Å^-1^ was covered (s = 4π sinθ/λ, where 2θ is the scattering angle). Data were analyzed using the ATSAS 2.8 package (37). The SEC-SAXS data were analyzed with CHROMISX and the forward scattering I(0) and the radius of gyration R_g_ were extracted from the Guinier approximation calculated with the AutoRG function within PRIMUS (38). GNOM (39) provided the pair distribution function P(r) of the particle, the maximum size D_max_ and the Porod volume. *Ab initio* reconstructions were generated with DAMMIF (40). Forty independent DAMMIF runs were compared and clustered into five main classes using DAMCLUST (41,42). Models within each class were superimposed by SUPCOMB (43) and averaged using DAMAVER (44). Structures were visualized using UCSF Chimera (30).

### Integrated modelling

The SAXS envelopes of SFTSV L protein were visualized using molmap within UCSF Chimera (30) at 15 Å resolution and the SFTSV L protein 3D EM map A was low-pass filtered to 5 Å to smoothen the volume. After filtering, the flexible C-terminal region became visible in the 3D EM map A, which was subsequently used to guide the initial fitting into the SAXS envelope, and further refined with the “fit in map” function of Chimera. The atomic model for SFTSV apo-L was fit into the 3D EM map A.

## RESULTS AND DISCUSSION

### Structure determination of SFTSV L protein by cryo-EM

We used a full-length construct of SFTSV L gene (strain AH12, accession HQ116417) for baculovirus-driven protein expression in insect cells. The L protein was either expressed as wild-type protein or as single-site mutants D112A (endonuclease inactive) or D1126A (RdRp inactive). The protein was purified to homogeneity, with a final yield of up to 11 mg pure SFTSV L protein per litre of cell culture, and used for cryo-EM experiments. Based on the single particle 3D reconstruction with a resolution of 3.8 Å (Supplementary Figure S1, Supplementary Table S2), the apo-SFTSV L protein (apo-L) structure was determined for ∼1100 residues mainly belonging to the RdRp core region (Figure 1). Density for the endonuclease domain was visible but the resolution was insufficient for *de novo* model building indicating mobility of this domain in the apo-conformation of the L protein. The apo-L structure determined by cryo-EM also lacks the C-terminal 500 residues. This domain seems to be very flexible, similar to the case of apo-LACV L, where the C-terminal 500 residues could also not be resolved using cryo-EM (12). Repeated classification resulted in an additional EM map that displayed additional electron density for the C-terminal domain (Supplementary Figure S2). However, the overall resolution was not high enough to confidently build additional parts of the protein compared to map A. In summary, we present a model for a substantial part of the SFTSV L protein in the apo-configuration and the cryo-EM map indicates the location and size of the C-terminal region.

**Figure 1.**
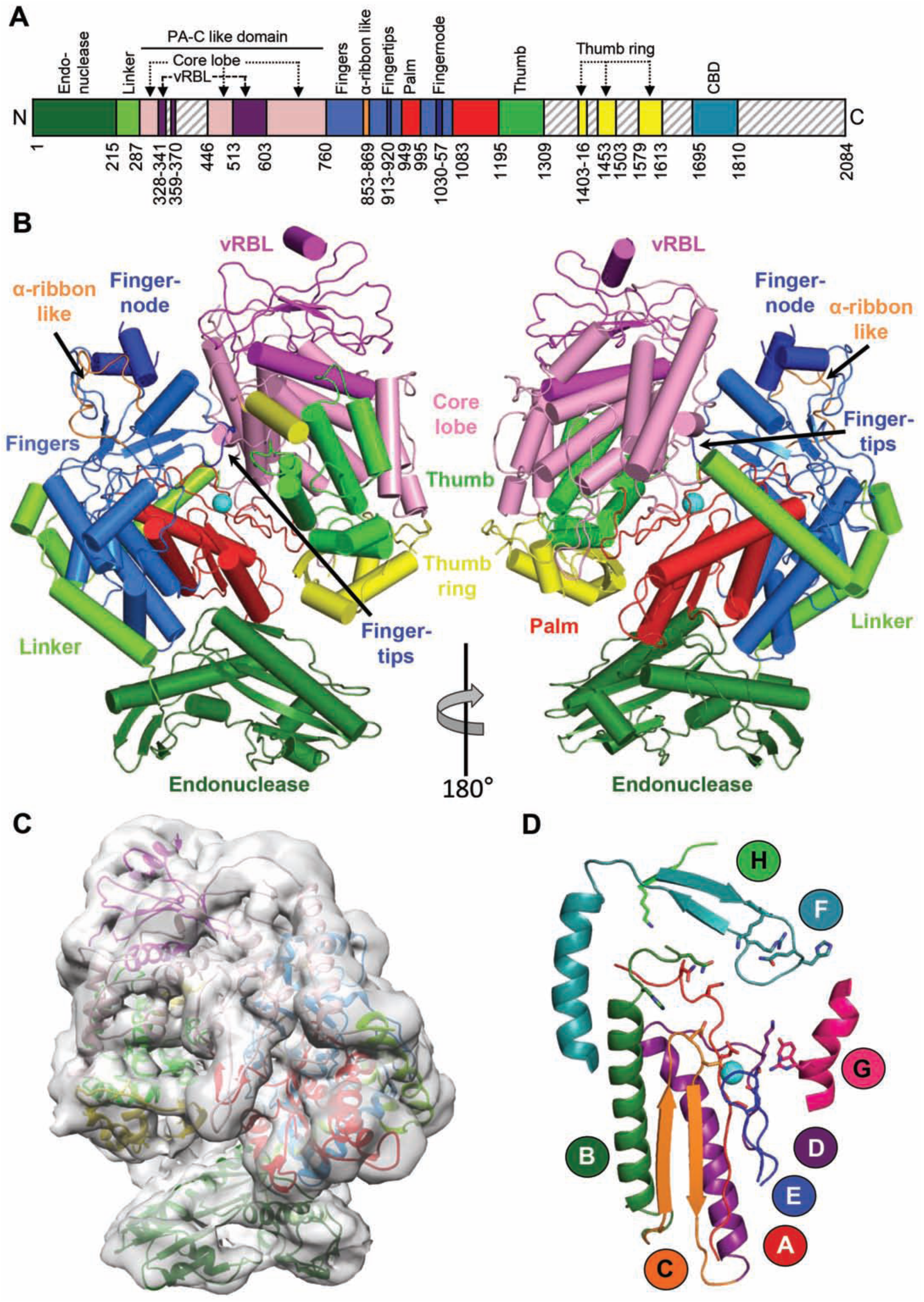
Apo structure of the SFTSV L protein. (**A**) Schematic linear representation of the domain structure of the SFTSV L protein. (**B**) Illustrated representation of two views of the apo-L cryo-EM structure as a ribbon diagram (PDB 6Y6K). Domains are colored as in (A). Magnesium ions in the active site are shown as spheres. A more detailed view and comparison with the LACV L (PDB 5AMQ) and the influenza virus polymerase complex (PDB 6QCV) are shown in Supplementary Figures S2, S3 and S4. (**C**) Superposition of the 5Å low-pass filtered cryo-EM map with the SFTSV apo-L structure model. (**D**) Representation of the polymerase core of the SFTSV L with the conserved motifs A-H as a ribbon diagram. Divalent magnesium ions are shown as spheres.

### SFTSV L protein is similar to the polymerase proteins of other segmented negative strand RNA viruses

SFTSV apo-L is structurally related to LACV L and the heterotrimeric influenza virus polymerase complex, although the similarity to LACV is higher (Supplementary Figures S2, S3 and S4). This is consistent with the phylogenetic relation between these viruses, which all belong to the sNSV group, and the previously described structural similarity between influenza virus polymerase and LACV L protein (12). Although the cryo-EM maps of SFTSV apo-L did not contain density sufficiently resolved to build the endonuclease domain *de novo*, it was possible to unambiguously place the SFTSV endonuclease structure recently obtained by X-ray crystallography (23) (residues 1-214 of PDB 6NTV) into the map by rigid body fitting (Figure 1C). The endonuclease domain (residues 1-214) is connected to an influenza virus PA-C like domain via an extended linker (residues 215-286) which wraps around the fingers and palm domain (Figure 1, Supplementary Alignment File). This linker is slightly shorter (∼15 residues) than in LACV L and has a more extended conformation (Supplementary Figures S2, S3 and S4). Similar to LACV, the PA-C like domain (residues 287-758, with residues 342-358 and 371-445 missing interpretable density) is divided into two lobes, an α-helical “core lobe” buttressing the palm and thumb domains of the L protein as well as an equivalent to the LACV vRNA binding lobe (vRBL) with a central β-sheet and at least two α-helices packing against both sides of the sheet (Figure 1 and Supplementary Figures S2, S3 and S4). A substantial part of this vRBL domain, which constitutes a 3’ vRNA binding site in LACV L protein, is missing from the SFTSV structure as it lacks defined density in the map. According to LACV L protein and influenza virus polymerase complex, we expect these unresolved regions to represent the “clamp”, involved in 3’ vRNA binding, and the “arch”, making contacts to the 5’ vRNA (Supplementary Figures S2, S3 and S4). The arch is also absent in the LACV L structure. The fingers, palm and thumb domains which contain most of the conserved RdRp active site motifs, are downstream of the PA-C like domain (Figure 1D, Supplementary Alignment File). The fingers domain (residues 759-948 and 995-1083) is again highly similar to the LACV counterpart (Supplementary Figures S3 and S4). However, the ribbon insertion (residues 853-870) extending from the fingers domain, is structurally different compared to LACV α-ribbon and influenza virus β-ribbon. In SFTSV L, the ribbon insertion is ∼40 residues shorter than in LACV L and structurally disordered (Figure 1B, Supplementary Figures S2, S3 and S4). The corresponding β-ribbon of influenza virus, which is involved in binding of the RNA duplex region of the promoter and contains the nuclear localization sequence, is ∼16 residues longer compared to SFTSV (Supplementary Figures S2, S3 and S4). The next specific feature of SFTSV L is the fingertips insertion (residues 913-920), a loop which is mostly conserved among bunyaviruses (45) and contains parts of the RdRp active site motif F (Figure 1D, Supplementary Alignment File). This loop has a different conformation in SFTSV compared to LACV, which is probably related to a conformational change induced by the 5’ RNA binding that binds as a stem-loop structure (the so-called ‘hook’) to the LACV L protein (45). A second insertion into the fingers domain is the so-called fingernode (residues 1030-1057), which is composed of two α-helices and involved in 5’ vRNA binding in the LACV and influenza virus structures. A small part of what is presumably the connecting loop between the α-helices is absent in the SFTSV apo-L structure (residues 1039-1045). The fingernode is structurally very similar to the LACV fingernode but different from its counterpart in influenza virus which has a β-hairpin insertion (Supplementary Figures S2, S3 and S4). The fingers domain is followed by the palm domain of the RdRp (residues 949-994 and 1084-1194), which is structurally highly similar to LACV and influenza virus (Figure 1, Supplementary Figures S2, S3 and S4). The palm domain contains the RdRp active site motifs A and C that are involved in the coordination of the divalent metal ions and the catalysis. Additionally, the active site motifs D and E are part of the palm domain (Figure 1D). In contrast to LACV palm domain, SFTSV L does not possess the so-called California insertion, whose function is unknown (Supplementary Figures S2, S3 and S4). The thumb domain of the RdRp (residues 1195-1308) is composed of a helical bundle and in contact with the previously mentioned PA-C like domain and the vRBL (Figure 1). Residues 1309-1402, which by analogy to the LACV and influenza virus structures presumably compose the priming loop and the bridge domain, are not visible in the SFTSV model (Figure 1, Supplementary Figures S2, S3 and S4). We cannot conclude on the length of the priming loop as the sequence identity to LACV is very low for this region (Supplementary Alignment File) and connected parts of the bridge are missing as well (Supplementary Figure S3). The so-called thumb ring is also only partly visible in the apo-L structure of SFTSV (visible parts include residues 1403-1416, 1452-1502, 1579-1587, and 1596-1612). Further regions missing from the cryo-EM structure are the lid and C-terminal domain including the CBD (Figure 1, Supplementary Figures S2, S3 and S4, Supplementary Alignment File). Overall, the SFTSV L protein has a very similar architecture as LACV and influenza virus polymerase proteins with some particular differences whose functional relevance will be interesting to determine. Regions missing from the SFTSV L and partly also from the LACV L structure indicate a high degree of flexibility of the C-terminal regions, especially the thumb ring, bridge, and C-terminal domain. Flexibility in the vRBL domain is most likely due to the absence of viral RNA to stabilize the structure but further structural data are needed to clarify this.

### 3’ and 5’ promoter RNA binding to SFTSV L protein

An important feature of all sNSV polymerase proteins is the ability to bind to the conserved RNA promoter ends, the complementary 3’ and 5’ termini of the genome segments. For LACV and influenza virus polymerases, distinct 3’ and 5’ RNA binding sites have been described (12,17). In electrophoretic mobility shift assays, we were able to detect interaction of SFTSV L protein with the conserved termini of all three genomic RNA segments (Figures 2A and B, Supplementary Figure S5). The migration behaviour of the protein–RNA complex in the native gel was different depending on whether 3’ or 5’ termini were bound (compare Figure 2A right and left panel) indicating that 3’ and 5’ RNAs induce distinct conformational states of the L protein. The migration behaviour of the complexes is consistent with previous findings on the LASV L protein (13). Interestingly, for the S segment, two different sequences of the conserved termini are published which differ from each other by an insertion/deletion of an A at position 9, counted from the 5’ end (Figure 2B). We found that both versions bind to the L protein with comparable affinities (Supplementary Figure S5).

**Figure 2.**
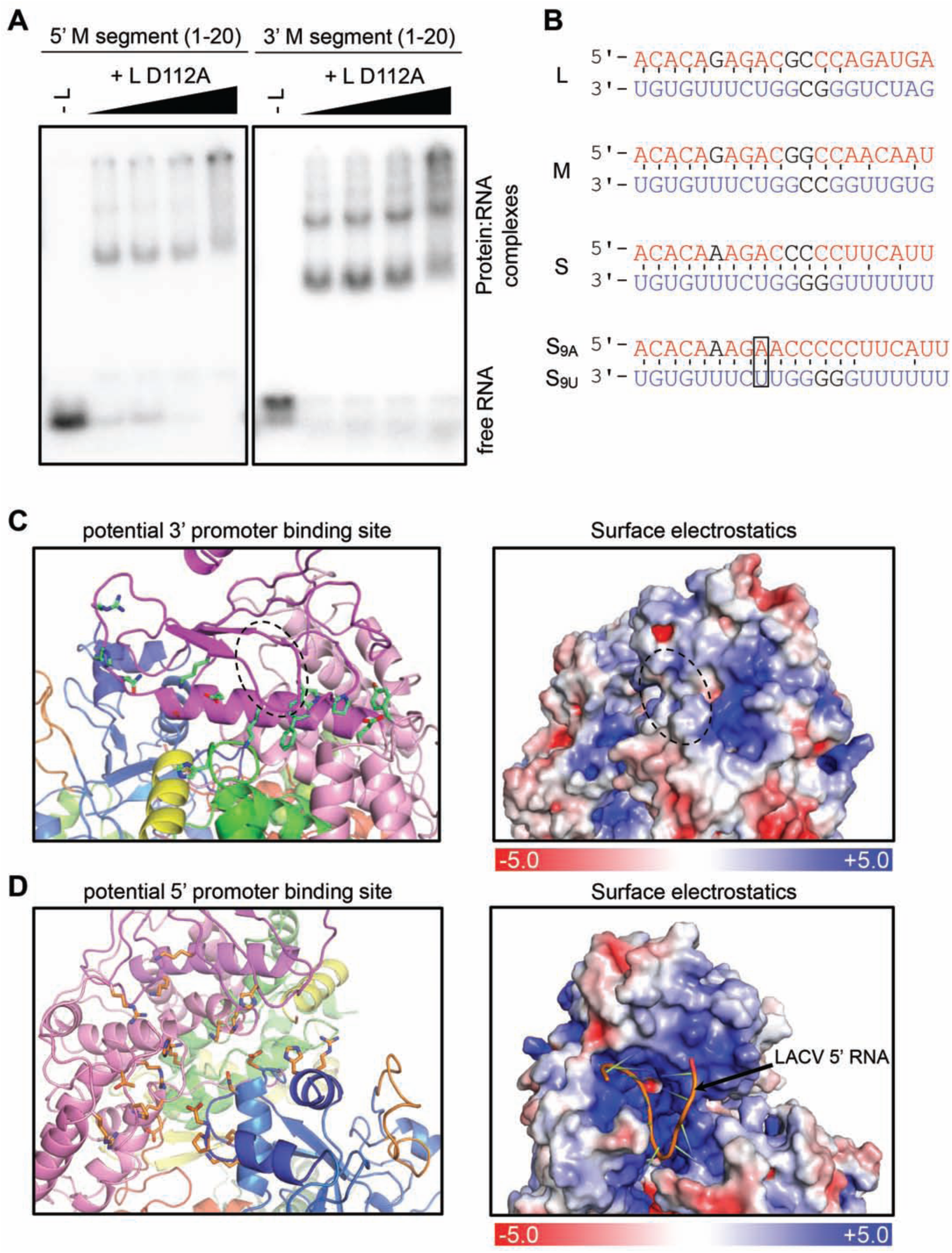
Interaction of the SFTSV L protein with its promoter RNAs. (**A**) Binding of SFTSV L protein to the 5’ and 3’ promoter ends (20 nt) of the M segment (Supplementary Table S1) was determined by an electrophoretic mobility shift assay. Increasing amounts of L protein (0 to 1.4 μM) were incubated with 0.2 μM of the indicated RNA (Supplementary Table S1). The protein-RNA complex was separated from the free RNA by native PAGE and visualized by phosphor screen autoradiography using a Typhoon scanner (GE Healthcare). (**B**) Conserved 3’ (red) and 5’ (blue) terminal sequences of the L, M and S segments. Watson-Crick base pairing is indicated by black lines. Bases which differ between, but are conserved within the segments are shown in black. Additional bases at position 9, found in some sequences (5’ S9A, 3’ S9U), are marked with a frame. Potential 3’ (**C**) and 5’ (**D**) promoter RNA binding sites are shown as cartoon representation (left). Domains are colored according to Figure 1. Residues potentially involved in protein-RNA interaction are shown as green (C) or orange (D) sticks. See also Supplementary Alignment File. Right panels display surface electrostatics of the potential 3’ and 5’ RNA binding sites generated by APBS within PyMol. LACV 5’ vRNA is shown as a cartoon within the SFTSV potential 5’ vRNA binding site. For comparison, analogous figures for LACV vRNA binding sites are presented in Supplementary Figure S6.

In LACV L protein, the 3’ RNA was bound in a narrow cleft leading away from the polymerase active site and formed by the PA-C like domain, the thumb and thumb ring with the clamp of the vRBL serving as a lid over the cleft (Supplementary Figure S6A). The 5’ vRNA was found to occupy a separate binding pocket in a hook-like conformation both in LACV as well as in influenza virus (12,17,18). By comparing with LACV L protein bound to the 3’ and 5’ promoter RNA, we suggest that the RNA binding sites of SFTSV L protein are in the equivalent locations. The 3’ RNA is likely bound in a positively charged cleft formed by the thumb and the PA-C like domain, especially the vRBL (Figure 2C). The vRBL domain, however, would have to undergo a slight conformational change as in the current conformation a loop between two β-strands (indicated in Figure 2C by a dashed circle) would block parts of the cleft. The clamp, which is missing clear density in the SFTSV cryo-EM map indicating flexibility in the apo-L could close the 3’ RNA-binding cleft on the top as observed for LACV (12). Although the clamp is missing we observed potential RNA interacting residues whose location and number in this area would correspond to LACV 3’ RNA binding site (compare Figure 2C and Supplementary Figure S6A, Supplementary Alignment File).

The 5’ hook RNA binding site in LACV is formed by the PA-C like and the fingers domains. We identified potential 5’ RNA-binding residues in SFTSV that closely match in location and number with the residues detected in 5’ hook binding in LACV (Figure 2D, Supplementary Figure S6B, Supplementary Alignment File). Similar to LACV we observe clustering of positively charged residues in this pocket that, in terms of size, allows to accommodate the 5’ RNA of LACV (Figure 2D, Supplementary Figure S6B). However, the exact residues interacting with the 5’ RNA cannot be predicted, as the 3D structure of the SFTSV hook remains unclear. In an attempt to characterize the possible conformations of the SFTSV 5’ RNA, we used the RNA secondary structure prediction program Mfold (46). Whereas in influenza virus and LACV the 5’ hook structure forms within the first 10 nucleotides, this seems rather unlikely for SFTSV 5’ termini judging from the predictions (Supplementary Figure S7). The prediction results in several possible hook structures but without consistency between the S, M and L segment promoters. It may be that the RNA hook structure is stabilized by base-specific protein–RNA interactions rather than base-pairing. However, the sequences reported for SFTSV genome segment termini vary significantly between the L and M segments on the one hand and the S segment on the other hand (Supplementary Figure S8). Therefore, additional structural information will be necessary for reliable conclusions about the putative 5’ hook structure and how it binds to the L protein. In summary, we observe potential 5’ and 3’ RNA binding sites in the SFTSV apo-L analogous to RNA binding sites reported for LACV L and also influenza virus polymerase.

### The SFTSV L protein is an active polymerase

To characterize the enzymatic properties of SFTSV L protein, we performed *in vitro* RNA synthesis based on the assay conditions established for LASV (13) using a highly purified SFTSV L protein (Figure 3A). To avoid RNA degradation during the enzymatic reactions we used an endonuclease active site mutant (D112A) of the L protein. SFTSV L protein produces a ∼35 nt RNA product independent of any primers, which was visualized by autoradiography based on incorporated radiolabelled α^32^P-GTP after denaturing PAGE (Figure 3B). An L protein carrying a mutation in the RdRp active site motif C (D1126A), which served as a negative control to demonstrate the specificity of the assay, was inactive. The SFTSV L protein RdRp was only active when both the 3’ template vRNA and the 5’ vRNA were present in the reaction (Figure 3B). The activating role of the 5’ end is well known for influenza polymerase, and has also been described for arenaviruses, although the proposed fold of the arenavirus 5’ hook has yet to be confirmed by a structure (13,16-18).

**Figure 3.**
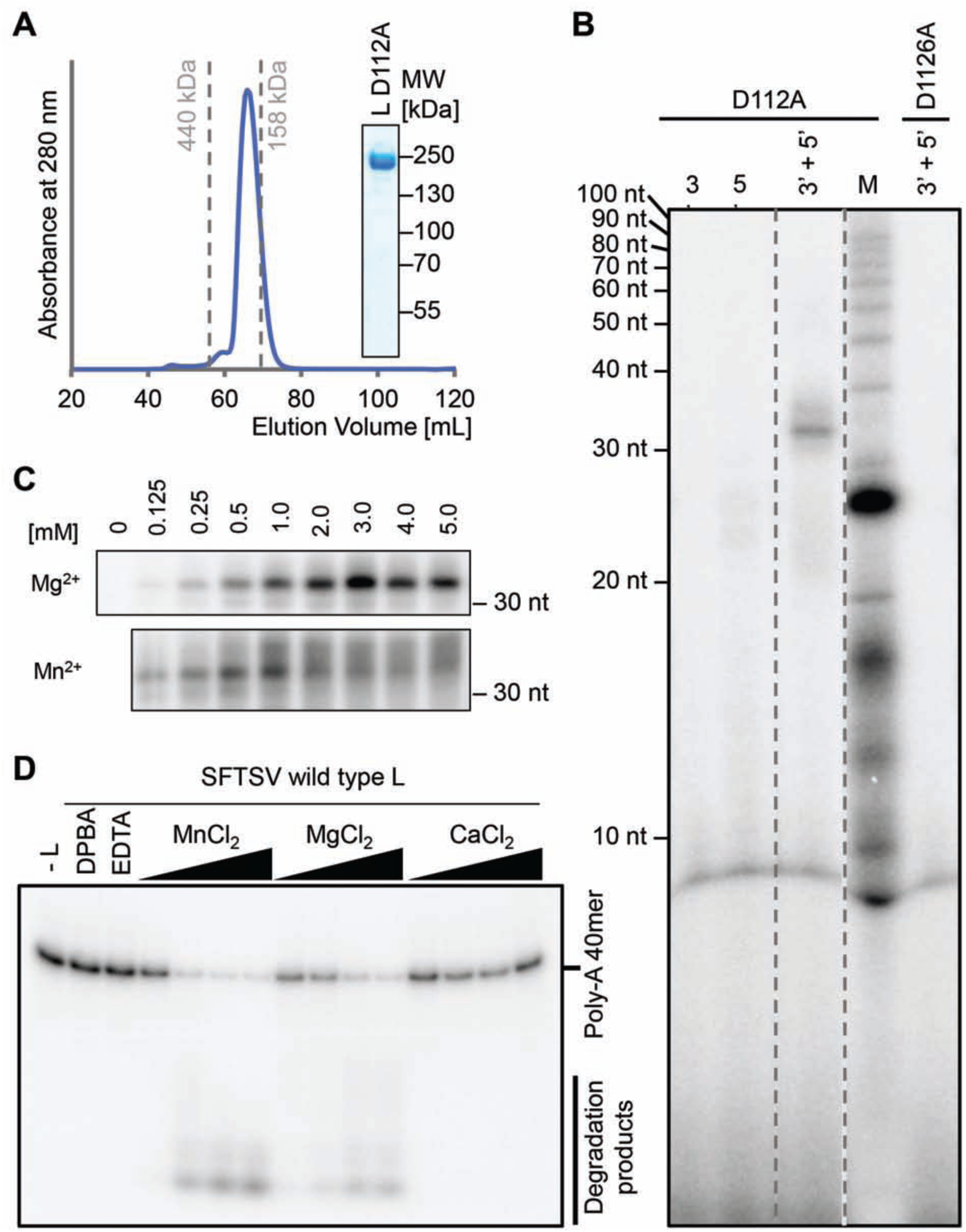
In vitro enzymatic activities of the SFTSV L protein. (**A**) Size exclusion chromatography and Coomassie stained SDS-PAGE analysis of purified SFTSV L (D112A) protein display the high purity and monodispersity of the L protein. Elution volumes of standard proteins (with sizes of 440 kDa and 158 kDa) for column calibration are indicated. (**B**) The SFTSV L protein mediates RNA synthesis in an *in vitro* polymerase assay. SFTSV L (D112A) or an RdRp catalytically inactive mutant (D1126A) were incubated with the conserved 5’ or/and 3’ terminal 20 nt of the M segment (Supplementary Table S1) in the presence of NTPs supplemented with [α]^32^P-GTP for 60 min at 30°C. RNA products were separated by denaturing gel electrophoresis and visualized by autoradiography. (**C**) RdRp activity of 500 nM SFTSV L (D112A) protein was analyzed in the presence of the indicated concentrations of MgCl_2_ or MnCl_2_. (**D**) 250 nM of wild-type L was incubated with ∼0.3 μM of radioactively labeled PolyA40 RNA substrate (Supplementary Table S1) in the presence of 5, 10, 25 and 50 µM of the indicated Me(II) at 37°C for 30 min. Reactions without L protein, EDTA or the known endonuclease-specific inhibitor DPBA were used as negative controls. Reaction products were separated on a denaturing polyacrylamide gel and visualized by autoradiography.

Notably, for the sequences of the conserved genome segment termini, particularly the S segment, different sequences have been published. For the S segment these sequences differ in the insertion/deletion of an A at position 9, counted from the 5’ terminus, or a U in case of the 3’ terminus (47-49). The S segment RNA including the A at position 9 (denoted as 9A) resulted in much stronger polymerase activity of the L protein compared to the S-segment promoter lacking the 9A, even though the affinity to the L protein seems to be comparable between the two S segment promoters (Supplementary Figures S5 and S9). The reduced polymerase activity with the S segment terminus lacking the 9A is consistent with results from Brennan *et al.* (2015) reporting that it was only possible to establish a reverse genetics system based on SFTSV S segment if the A at position 9 of the 5’ end as well as a corresponding U at the complementary 3’ end were present (49). Even though, we cannot conclude on the role of this 9A residue from our apo-L structure, this should be noted for future studies.

As already described for other polymerases (50,51), enzymatic activity was also dependent on divalent metal ions. SFTSV RdRp displayed strong activity in the presence of magnesium ions with an activity plateau reached at Mg^2+^ concentrations of >2 mM (Figure 3C), which is similar to what has been reported for LASV RdRp for manganese ion concentrations (13). The presence of 5 mM Mg^2+^, nucleotides and both 3’ and 5’ promoter RNA was defined as the standard reaction conditions for *de novo* replication by SFTSV L protein and resulted in a single and strong product band after 60 min incubation at 30°C (Figure 3B). In presence of manganese ions, the RdRp activity was also detectable. However, with higher Mn^2+^ concentrations (>1 mM) the product band got more diffuse resulting either from digestion by the endonuclease (stimulated by the high concentration of Mn^2+^) or from less accurate replication initiation by the RdRp. For arenaviruses, the RdRp activity was greatly enhanced when the 5’ end had a single nucleotide G-overhang compared to the complementary 3’ promoter template strand (13,16), originating from a prime-and-realign mechanism for genome replication. The reason for this enhancing effect was speculated to result from either improved promoter binding or effects on the secondary structure of the 5’ hook. However, such an enhancing effect was not observed for SFTSV L protein (Supplementary Figure S10A).

The product detected in our *in vitro* primer-independent genome replication reactions was larger than expected. This has previously been observed for LASV in i*n* vitro polymerase assays (13). It was argued that this could be due to missing termination signals as the assay contains only physically separated short promoter strands of 20 nt rather than the continuous RNA genome comprising all necessary cis-acting signals (13). Another possibility is that the template and the completely complementary product form very stable RNA duplexes, which are not separated even by the denaturing PAGE conditions used. We tested this hypothesis using 20 nt perfectly complementary RNA which was fluorescently labelled instead of radioactively labelled, and performed denaturing PAGE at either 20°C (as usually done) or higher temperatures (60°C). Indeed, we were unable to separate the RNA duplex at electrophoresis temperatures of 20°C even though samples were heated to 95°C and supplemented with denaturing loading buffer prior to denaturing PAGE, whereas at 60°C we detected the RNA at the expected height (Supplementary Figure S10B). Therefore, we can explain the large size of our products in the assays by the difficulty to separate perfectly complementary RNA, a product that cannot be avoided when investigating viral genome replication. However, this does not compromise the specificity of our assay.

In summary, we established a polymerase assay for the SFTSV L protein in which the L protein synthesizes a specific product with the minimal components of Mg^2+^, nucleotides, conserved 3’ template and 5’ promoter strand.

### L protein of SFTSV contains an active endonuclease

We tested the SFTSV full-length L protein for endonuclease activity using a ribonuclease assay with a radiolabeled 40mer ssRNA substrate. Substrate degradation in the presence of different divalent metal ions was detected by denaturing PAGE and autoradiography. We found that the endonuclease in SFTSV L protein was active in presence of either manganese or magnesium ions but not calcium ions (Figure 3D). Residual activity was also detected when zinc, nickel and cobalt ions were added to the reaction (Supplementary Figure S10C). These results do not entirely match with reports for the isolated endonuclease domains of SFTSV and closely related Toscana virus (TOSV), for which the endonuclease was inactive in the presence of magnesium ions (23,52). As a negative control, we expressed a full-length L protein with a mutation in the endonuclease active site (D112A). Contrary to previous findings on the isolated endonuclease domains of SFTSV and TOSV with mutations of the active site (23,52), the full-length D112A L protein mutant showed some residual RNA degradation activity in the endonuclease assay. We hence added either EDTA or the known endonuclease-specific inhibitor DPBA (8) to our negative controls (Figure 3D).

In conclusion, we demonstrate that the endonuclease activity of the full-length SFTSV L protein is dependent on divalent manganese or magnesium cations. As in our polymerase and protein-RNA interaction assays we also observed some degradation of the polymerase products and promoter RNA, we conclude that the endonuclease cleaves both viral and non-viral RNA. This suggests that the endonuclease activity is not sequence-specific and underlines the need for activity regulation on the one hand and the need for protection of viral RNA by binding to the L protein or viral nucleoprotein on the other hand.

### Investigation of the mechanism of genome replication initiation

Based mainly on sequencing data from a number of sNSV a prime-and-realign mechanism has been proposed for *de novo* initiation of genome replication (19-21) and possibly transcription (53-55). There is substantial evidence that the LASV L protein uses a prime-and-realign mechanism during replication initiation (13). LASV L protein initiates genome replication at position +2 of the template strand, produces a dinucleotide primer and realigns the 5’ end of this primer to positions -1 and +1 of the template resulting in a single nucleotide overhang of the product relative to the complementary 3’ template (13). To characterize the mechanism of genome replication initiation of the SFTSV L protein, we performed primer extension polymerase assays in the presence of different radioactively labelled RNA primers and compared the product size with the size of the *de novo* product (from polymerase reaction with radiolabeled α^32^P-GTP) (Figure 4A). In Figure 4C, we provide an overview of all possible products for each of the primers used. In all reactions, we observe only one single product band, indicating that the SFTSV L protein employs only one replication initiation site on the template (Figure 4A). We observe the same product size as the *de novo* product when using ACA or AC primers in the reaction. If CA and CAC primers are provided, the resulting products are ∼1 nt smaller than the *de novo* product (Figures 4A and B). Therefore, there are three possible scenarios for genome replication initiation: (1) replication is initiated terminally at position +1 of the template strand, (2) replication is initiated internally at position +3 and the nascent ACA primer is subsequently re-aligned to position -2/-1/+1 of the template strand, or (3) replication is initiated internally at position +3 and the nascent AC or ACA primer is re-aligned to the terminal position +1/+2/+3 of the template (Figure 4C). For further clarification, we used a longer primer (ACACAAA) for the reaction, which is complementary to the first seven nucleotides of the template strand and should only support terminal initiation (Figure 4C). This primer was incorporated in a product resulting in exactly the same size as the *de novo* product (Figures 4A and B) showing that we can exclude realignment of the primer to position -2/-1/+1. Although it is not possible to reliably discriminate between terminal initiation or priming and subsequent realigning of the primer to the terminal position of the template (Figure 4D), terminal initiation seems more likely as there is only one defined product band detected. In case of priming internally and realigning to the terminal position one would expect to also see a minor product band, two nucleotides shorter than the main product, resulting from missing realignment. At least, this has been observed for LASV L protein, which applies a prime-and-realign mechanism for genome replication initiation (13). We conclude that SFTSV initiates its genome replication on a vRNA template either terminally and without applying a prime-and-realign mechanism similar to influenza virus cRNA synthesis (22) or by priming at position +3 and subsequent realignment to the terminal position +1 (Figure 4D). In any case, SFTSV L protein does not seem to produce a single or di-nucleotide overhang of the 5’ promoter end compared to the 3’ end. A scenario proposed for hantavirus genome replication is that after internal priming and subsequent realignment a single-nucleotide overhang is removed by the endonuclease resulting in a monophosphorylated 5’ terminus (21). However, this mechanism is rather unlikely to occur during SFTSV replication initiation as we used an endonuclease inactive mutant of the L protein in our assays. Even though this mutant showed residual endonuclease activity, full cleavage would be necessary to produce the single product band we observed.

**Figure 4.**
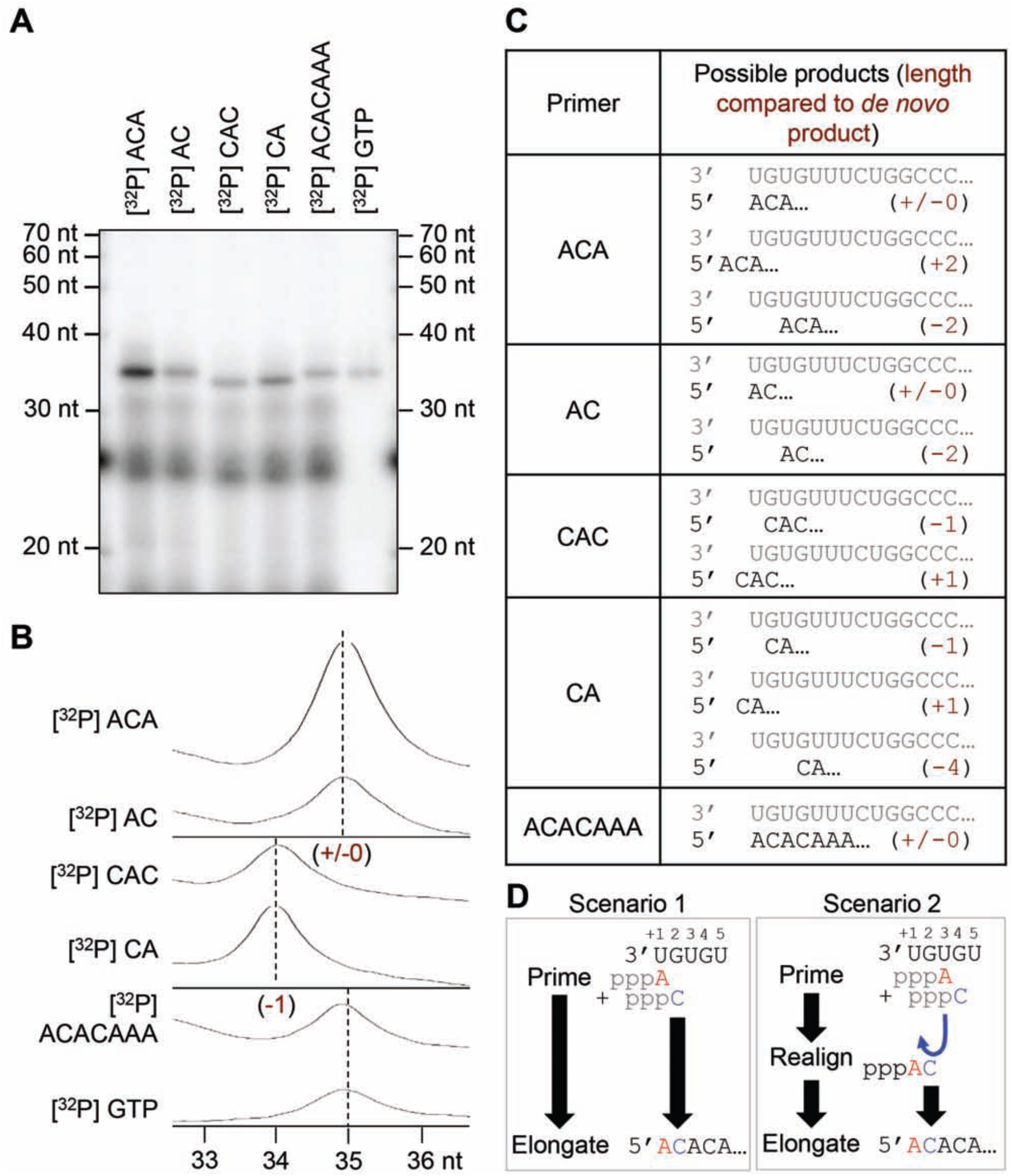
Initiation of replication. (**A**) RNA products synthesized by the SFTSV L (D112A) protein in the presence of the radioactively labeled primers listed in (C). The experiment was performed as described for the standard polymerase assay (see Materials and Methods). Where indicated, radioactively labeled primers were used instead of [α]^32^P-GTP. (**B**) The intensity profiles of the gel lanes from (A) were analyzed using ImageJ software (60) and illustrate the size differences between the product bands shown in (A). Sizes were determined by linear regression using the RNA marker (lower X-axis). (**C**) The table summarizes the RNA oligonucleotides used as primers in this experiment and the possible products with and without realignment. In brackets the length of the product RNA, relative to the *de novo* product is given. (**D**) Schematic representation of the possible priming scenarios. Scenario 1 depicts terminal initiation: ATP primes the reaction by binding to the first nucleotide (position +1) of the template and is further elongated without realignment, resulting in a product with the same length as the *de novo* reaction product (+/-0). Scenario 2 depicts internal initiation and realignment: the reaction is primed internally by binding of the first ATP to the position +3 and the addition of a C to form a di-nucleotide primer followed by dissociation of the AC dinucleotide and its realignment to position +1 and +2. This realigned AC dinucleotide is then elongated, resulting in the same product as the terminal initiation (+/-0).

### SFTSV L protein contains an active cap-binding domain

In analogy to influenza virus polymerase, the C-terminal region of the bunyavirus L protein has been suggested to contain the CBD that is needed for the cap-snatching mechanism employed by sNSV for transcription priming (7). Recently, the CBD of closely related RVFV has been determined and the residues interacting with a co-crystallized cap-analogue m^7^GTP have been proven to be essential for virus transcription in a cell-based minireplicon system (11). As the C-terminal domain of SFTSV apo-L could not be resolved from the cryo-EM data, we expressed only the putative CBD of SFTSV in *E. coli* (residues 1695-1810). The purified protein crystallized as a monomer in complex with an m^7^GTP cap-analogue and the crystals diffracted to 1.35 Å resolution (Supplementary Table S3). The SFTSV CBD is structurally very similar to RVFV CBD with a 7-stranded mixed β-sheet, a β-hairpin at the periphery of the domain and a long α-helix packed against the β-sheet (Figure 5A). The key residues responsible for the interaction with the m^7^GTP are functionally conserved among phenuiviruses (Figures 5B, Supplementary Alignment File). The m^7^GTP is stacked between the two aromatic side chains of F1703 and Y1719 extending from the first β-strand and the β-hairpin, respectively. Further interactions are observed between Q1707 from the hinge between the first β-strand and the β-hairpin, L1772 from the end of the long α-helix and the carbonyl group of D1771 (Supplementary Table S4, Supplementary Figure S11). We characterized the interaction of m^7^GTP and the protein by isothermal titration calorimetry (ITC) and thermal stability assays (Figures 5C and D). ITC data demonstrate specific interaction of SFTSV CBD with m^7^GTP cap-analogue in contrast to extremely weak interaction with unmethylated GTP (Figure 5D). Analysis of ITC data provided a dissociation constant of K_D_ ∼138 µM for m^7^GTP binding to SFTSV CBD which is 5-fold lower than the K_D_ observed for the RVFV domain but still quite high compared to influenza virus PB2 CBD (K_D_∼1.5 µM) or cellular cap-binding proteins (K_D_∼10-13 nM) (7,11,56-58). Thermal stability assays revealed a larger shift in the melting temperature (T_m_) for SFTSV CBD upon addition of m^7^GTP compared to GTP or ATP (Figure 5C). Consistent with the higher affinity of SFTSV CBD for m^7^GTP determined by ITC, the shift in T_m_ was also higher compared to RVFV CBD (compare Figure 5C and Supplementary Figure S12A): +8°C for SFTSV vs. +4.5°C for RVFV in the presence of 10 mM m^7^GTP. We observe this difference between SFTSV and RVFV in both assays even though the number of interactions between the protein and the m^7^GTP ligand is only slightly higher in SFTSV compared to RVFV (compare Supplementary Table S4 with data from Gogrefe *et al.*) (11). To provide additional evidence for the essential role of residues interacting with m^7^GTP, we expressed and purified mutants of the CBD - F1703A, Y1719A and Q1707A - and tested them in the thermal stability assay (Supplementary Figure S12B). As expected, the mutated CBDs were not significantly thermally stabilized in the presence of m^7^GTP and the T_m_ of the mutated proteins was ∼3-4°C lower than the T_m_ of wild-type CBD indicating overall lower stability of the domain upon single mutation.

**Figure 5.**
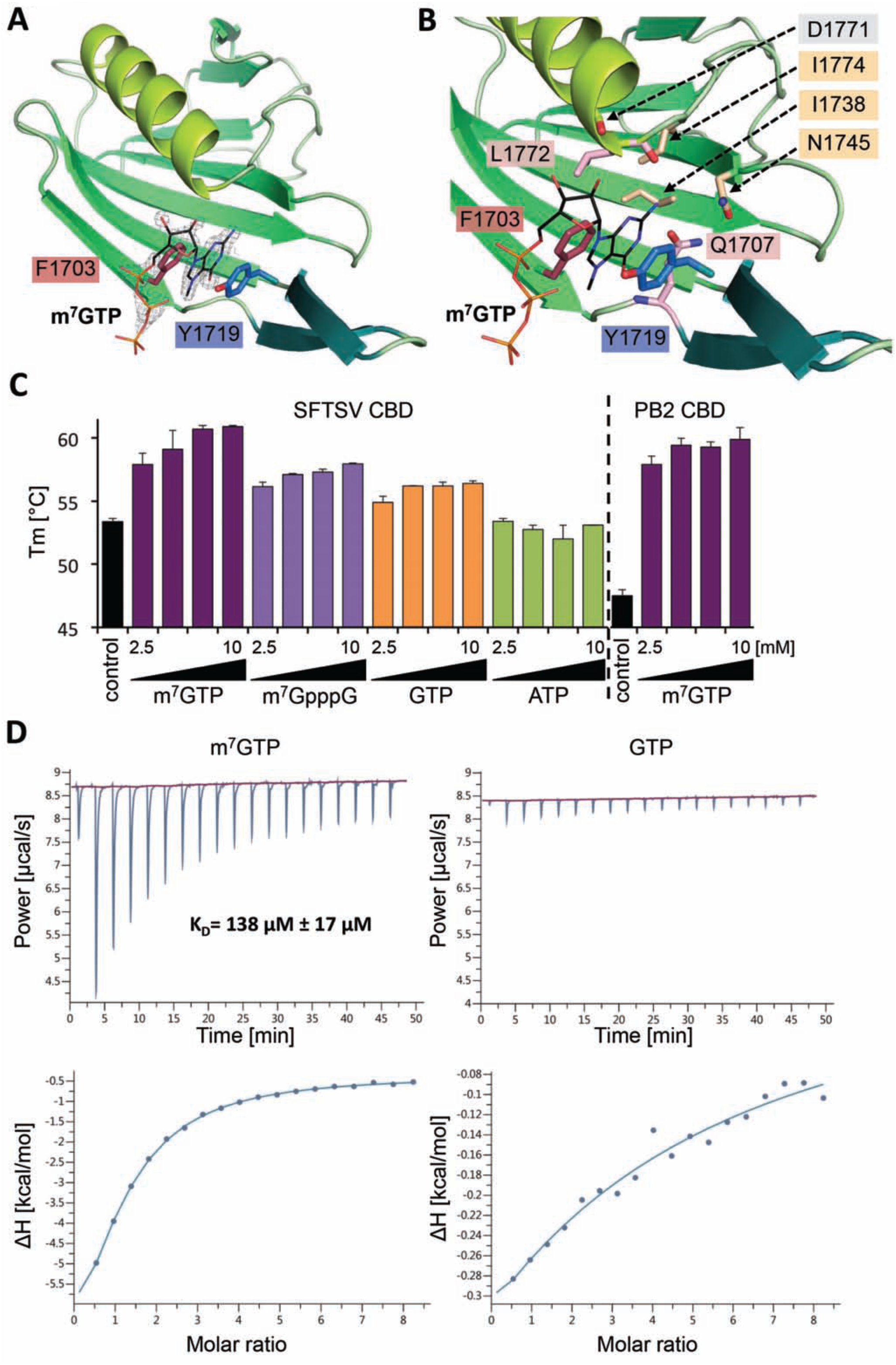
Structure of SFTSV CBD and m^7^GTP binding. (**A**) The figure shows SFTSV CBD crystal structure in complex with an m^7^GTP. SFTSV CBD is presented as a ribbon diagram with the side chains of the two aromatic residues (F1703, Y1719) involved in stacking interaction with the m^7^GTP ligand shown as sticks. m^7^GTP is presented as lines and the surrounding electron density (2|Fo|-|Fc| omit map at 2.0σ) as grey mesh. (**B**) In a close-up of the m^7^GTP binding site, the protein is shown as ribbon diagram and side chains involved in m^7^GTP binding as well as the carbonyl oxygen of residue D1771 are presented as sticks. The m^7^GTP is shown as lines. The residues I1774, I1738, N1745, are involved in stabilizing the binding site cavity, D1771 (carbonyl oxygen), Q1707 and L1772 directly interact with m^7^GTP. A detailed list of interactions between the CBD and the m^7^GTP ligand is given in Supplementary Table S4 and a ligand plot in Supplementary Figure S11. (**C**) Thermal stability of SFTSV CBD and influenza A virus PB2 CBD was tested in presence and absence of different concentrations (2.5, 5.0, 7.5, and 10 mM) of m^7^GTP, m^7^GpppG, GTP and ATP. Melting temperatures (T_m_) are presented as mean and standard deviations of three independent measurements (n=3). (**D**) The affinity of SFTSV CBD for m^7^GTP and GTP was measured by isothermal titration calorimetry at 25°C. A representative titration curve is shown. Titrations were done three times with 150 µM SFTSV CBD in the cell and 5.0-6.5 mM m^7^GTP or GTP in the syringe. The upper panel shows the raw data, the lower panel the integrated data fitted to a single-site binding model with the stoichiometry fixed to 1. The dissociation constant K_D_ is given as a mean and standard deviation of three independent measurements for m^7^GTP.

Although the cap-binding ability of the isolated domain seems to be clearly present we were unable to establish cap-dependent transcription assays for the full-length SFTSV L protein. In the presence of a 16 nt capped or uncapped primer, designed to complementarily align with 3 nucleotides of the 3’ end of the template strand, the L protein synthesized a product which was about ∼12-16 nt larger than the *de novo* product (Supplementary Figure S13A). This result indicates that the primer was incorporated into the final product, but independent of the need for a 5’ cap.

Binding of a capped primer to the CBD of the L protein could lead to endonuclease cleavage products of specific length depending on the distance between the CBD and the endonuclease active site. Therefore, we tested for cap-dependent endonuclease activity using poly-A RNA with either cap0 (m^7^GTP), cap1 (m^7^GpppNm) or no cap at the 5’ end but did not detect any specific cleavage product (Supplementary Figure S13B). In summary, we provided evidence for a functional CBD within SFTSV L protein (residues 1695-1810) structurally similar to RVFV and influenza virus CBD, but were unable to demonstrate any cap-dependent polymerase activity of the full-length L protein. It remains unclear what activates the cap-binding function of the full-length L protein. Interaction with host factors or the viral nucleoprotein might be necessary for the L protein to switch to transcription mode (7). This is also conceivable regarding the comparably low affinity detected for m^7^GTP binding to the CBD *in vitro*. Similar results have been reported for RVFV CBD (11). Further studies are required to fully elucidate how bunyavirus cap snatching and cap-dependent transcription works.

### Integrative modelling

We used the pure, monodisperse and monomeric full-length SFTSV L protein to perform SAXS experiments and obtain a low-resolution structure of the L protein in solution (Figure 3A, Supplementary Figure S14A). Three representative SAXS models were obtained by clustering analysis of forty *ab initio* dummy atom models and averaging of the structures within each of the three biggest clusters. All three models feature a compact core domain with a hollow center that is decorated with at least one protruding sub-domain (Supplementary Figure S14B), as observed for LASV L protein previously (13). However, the three SAXS models differ slightly in the size and location of a second protrusion from the core domain. We used these SAXS models for integrative modelling of an SFTSV L protein containing the described incomplete SFTSV cryo-EM structure (Figure 6A, Supplementary Figure S15). The cryo-EM map overall fits the SAXS envelopes, to the exception of the volume of the endonuclease domain (Figure 6A, dashed circle) indicating mobility of this domain relative to the polymerase core in solution. As mentioned above, the endonuclease domain structure was added to the model by rigid-body fitting of the recently published crystal structure of the isolated domain (23). As our model of the L protein core region and the endonuclease crystal structure overlap within 14 residues, we were able to connect these two structures and include the endonuclease in our final model. These overlapping 14 residues form a helix, which has been demonstrated to be very flexible in its position relative to the endonuclease core, suggesting a role in regulation of the endonuclease activity by controlling access to the active site (23). The structural data presented here suggest classification of this flexible helix as the endonuclease-polymerase linker region (Figures 1A and B). In the SFTSV apo-L structure, this linker is in an extended conformation (Supplementary Figure S4A). However, it is conceivable that the linker can also be present in a more collapsed conformation as observed in LACV and influenza virus polymerase proteins, depending on the functional state of the L protein, which would be compatible with the hypothesized function in endonuclease activity control. Indeed, in the integrated model, it is conceivable that the endonuclease domain can rotate relative to the polymerase core and thereby fill the larger protrusion of the SAXS envelope (Figure 6B). In that state, the part of the endonuclease denoted as additional β-sheet in phenuivirus endonucleases, which was predicted to play a role in protein-protein interactions (23), could make contacts to the polymerase core or unresolved C-terminal region (Figure 6B).

Focussing on the C-terminal region of the L protein, in both the cryo-EM map as well as the SAXS envelopes, we observe low-resolution density volume into which it was not possible to build a structural model *de novo* (Figure 6A, indicated empty volume). This volume likely contains the C-terminal region especially missing parts of the thumb ring, bridge and lid. Notably, in the crystal structure of the arenavirus L protein C terminus the long linker connection of the CBD to the L protein probably enables high mobility of this domain (10). The second protrusion of the SAXS envelope, which is less pronounced and not visible in all structures, might correspond to the CBD. However, this remains purely speculative based on the data we have and we therefore did not include the CBD in the model.

**Figure 6.**
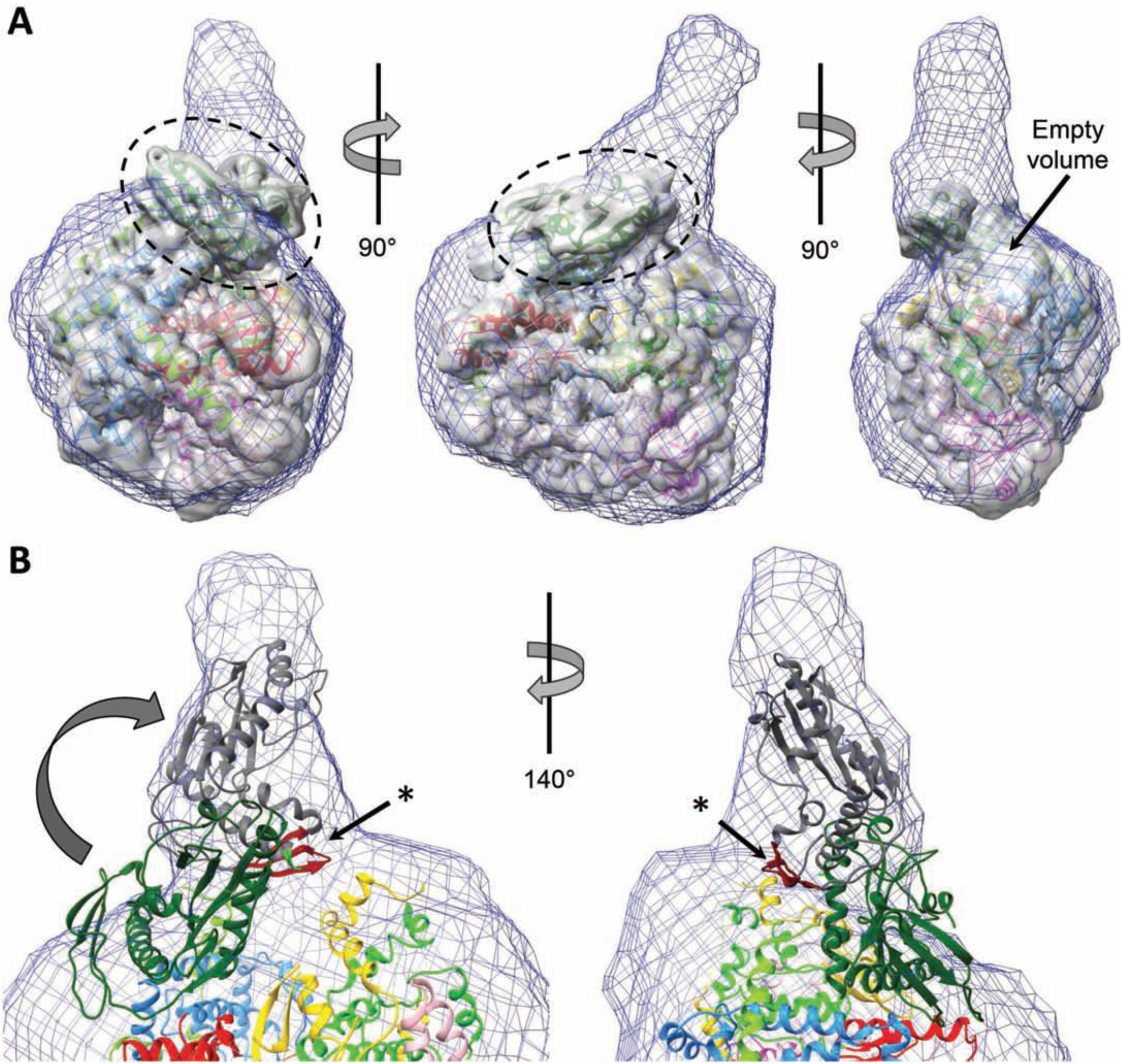
Integrative modelling of SFTSV apo-L protein. (**A**) A superposition of the SAXS envelope of cluster 1 (blue mesh), a 5Å low-pass filtered cryo-EM mapA (grey surface) and the structure model (ribbon diagram, colored according to Figure 1) is presented in three different orientations. The endonuclease domain, which is sticking out of the SAXS envelope, is marked by a dashed circle. An empty volume is also indicated. (**B**) Close-up of the endonuclease domain in the superposition of SAXS envelope and structure model. Potential movement of the endonuclease domain is indicated by an arrow and an alternative conformation depicted as grey ribbon diagram. An asterisk indicates the β-sheet (colored in red), for which a role in protein-protein interactions has been proposed.

Combining high- and low-resolution structural data from cryo-EM and SAXS, we showed that the conformation of the polymerase is globally similar in solution and in cryo-EM. Furthermore, it is conceivable that the endonuclease is able to rotate in respect to the polymerase core.

## CONCLUSIONS

Here we provide a comprehensive characterization of the SFTSV L protein structure and function using a combination of cryo-EM, X-ray crystallography, SAXS and biochemical assays. The structure of SFTSV L protein in the apo conformation closely resembles the LACV L protein and influenza virus polymerase complex structures and in analogy, it allows for the prediction of the RNA binding sites in SFTSV L protein. Notably, significant differences between these three viral polymerases include the length and conformation of the ribbon-like insertion, the position of the endonuclease domain relative to the polymerase core, and the conformation of the endonuclease linker region. In particular, the ribbon-like insertion has been speculated to be involved in L protein-nucleoprotein interactions, an interface that must be highly specific for each virus (12,18). Integrating structural data from SAXS and cryo-EM experiments, we concluded for flexibility of the endonuclease domain relative to the polymerase core region in solution. These observations fit to recently published analyses (23) and might explain the different positioning of the endonuclease relative to the polymerase core between LACV and SFTSV L structures. The C-terminal part of the bunyavirus L protein seems to be highly dynamic and could hence not be sufficiently resolved in the cryo-EM map. This has been previously observed for influenza virus PB2 (59) and LACV L (12). Mechanisms for stabilization of this region by viral RNA or other factors have to be defined in future studies of functionally relevant stages, *i.e.* initiation, elongation and termination of transcription. Additionally, the expression and purification procedures as well as the biochemical assays established will foster further structural and functional studies on bunyavirus L proteins. We demonstrated that the L protein of SFTSV binds to both 3’ and 5’ promoter RNA *in vitro* inducing distinct conformational stages as concluded from electrophoretic mobility shift experiments. Furthermore, we show that that SFTSV L likely initiates genome replication on vRNA *de novo* without applying a prime-and-realign mechanism and that cap-dependent transcription requires an unknown switch. Altogether, the structural and functional data presented here on the L protein of SFTSV improve our understanding of this complex and multidomain protein that is essential for viral replication. We also provide significant insights into the commonalities and differences between sNSV polymerase proteins, which will be particularly important for the development of broad-acting antivirals.

## Supporting information

Supplementary Alignment File

## DATA AVAILABILITY

Structural data are available from the PDB database (accession numbers 6Y6K and 6XYA). EM data have been deposited with the EMDB (accession number EMD-10706). All remaining relevant data are included in the manuscript and its supplementary material.

## SUPPLEMENTARY DATA

Supplementary Data are available at NAR online.

## ACKNOWLEDGEMENTS

The synchrotron MX data were collected at beamline P14 operated by EMBL Hamburg at the PETRA III storage ring (DESY, Hamburg, Germany). We would like to thank Thomas Schneider and Isabel Bento for the assistance in using the beamline and Isabel Bento additionally for helpful advice. The synchrotron SAXS data were collected at beamline P12 operated by EMBL Hamburg at the PETRA III storage ring (DESY, Hamburg, Germany). We would like to thank Tobias Graewert, Haydyn Mertens and Cy Jeffries for the assistance in using the beamline. We also thank the team of the EMBL Hamburg Sample Preparation and Crystallization (SPC) facility for support of ITC measurements. The authors thank Imre Berger for providing the DH10EMBacY *E. coli* and the team of the Eukaryotic Expression Facility (EEF) at EMBL Grenoble for support and advice.

