## Supplementary Figure, Supplementary Table for "Structural and functional characterization of the Severe fever with thrombocytopenia syndrome virus L protein"

#### Supplementary Tables

**Supplementary Table S1. Synthetic RNA used in the assays**

The table lists the RNA oligonucleotides that were used in the described assays. The specific sequence, length in nucleotides and the identifier used to label the RNA in the experimental descriptions are given. RNAs were synthesized by Biomers.

**Supplementary Table S2. Cryo-EM data collection, refinement and validation statistics.**

This table provides the statistics for the data collection, refinement and structure validation of the cryo-EM structure.

**Supplementary Table S3 Crystallographic table.**

This table provides the statistics for the data collection, refinement and structure validation of the crystal structure.

**Supplementary Table S4. Interactions with $m^7$GTP table**

This table lists the interactions between SFTSV CBD and the co-crystallized $m^7$GTP cap-analogue as generated by PDBsum (1).

#### Supplementary Figures

**Supplementary Figure S1: Cryo-EM of SFTSV L protein.**

(**A**) Representative micrograph of SFTSV L protein in vitreous ice after MotionCorr2 at defocus of ~3 µm. **(B)** 2D class averages of the SFTSV L protein. Front view: **(C)** Angular distribution for particle projections included in the final 3D reconstruction visualized as a sphere around the electron density map. **(D)** Electron density map (left) and colored according to the local resolution (right). Back view: **(E)** Angular distribution for particle projections included in the final 3D reconstruction visualized as a sphere around the electron density map. **(F)** Electron density map (left) and colored according to the local resolution (right). **(G)** Fourrier Shell Correlaton (FSC) curves for the SFTSV L protein between the two independently refined half-maps shows the overall resolution of the electron density map. (**H**) Close-up on a superimposition of the map (displayed as Fo-Fc map at 3σ) and the model in the core region of the map.

**Supplementary Figure S2. Comparison of the linear domain arrangement between SFTSV L, LACV L and influenza virus polymerase complex.**

Schematic linear representation of the domain structure of the SFTSV L protein (PDB 6Y6K) (top) from N to C terminus in comparison to LACV L protein (PDB 5AMQ, middle) and heterotrimeric (PA-PB1-PB2) influenza polymerase complex (PDB 6QCV, bottom). Structurally and/or functionally equivalent domains have similar colors. Abbreviations used as follows: ext. – extension; ins. – insertion. The residue numbers are given in respect to the respective structure. Regions larger than 10 amino acids not included in the respective models are colored with grey stripes.

**Supplementary Figure S3. Overall structure comparison between SFTSV L, LACV L and influenza virus polymerase complex.**

(**A**) Front and (**B**) back view of the structures of SFTSV apo-L (PDB 6Y6K), LACV L (PDB 5AMQ) and the heterotrimeric (PA-PB1-PB2) influenza virus polymerase complex (PDB 6QCV) (from left to right). Protein domains are colored and labelled according to Figure 1 and Supplementary Figure S2.

**Supplementary Figure S4. Detailed structure comparison between SFTSV L, LACV L and influenza virus polymerase complex.**

For simpler comparison, parts of structures of SFTSV apo-L (PDB 6Y6K), LACV L (PDB 5AMQ) and the heterotrimeric (PA-PB1-PB2) influenza virus polymerase complex (PDB 6QCV) are shown side by side: (**A**) regions corresponding to influenza virus PA, (**B**) PB1-like regions, and (**C**) PB2-like regions. Colors correspond to Supplementary Figures S2 and S3.

**Supplementary Figure S5. Binding of the conserved vRNA promoter ends to the SFTSV L protein.**

Binding of SFTS L (D112A) protein to the 5' (**A**) and 3' ends (**B**) (20nt) of the L, M, and S segments (Supplementary Table S1) was determined by an electrophoretic mobility shift assay. Increasing concentrations of L protein (0 to 1.4 µM) were incubated with 0.2 µM of the indicated RNA (Supplementary Table S1). The protein-RNA complex was separated from the free RNA by native PAGE and visualized by phosphor screen autoradiography using a Typhoon scanner (GE Healthcare).

**Supplementary Figure S6: vRNA binding sites of the LACV L protein.**

3' (**A**) and 5' (**B**) promoter RNA binding sites of the LACV L protein (PDB 5AMQ) shown as cartoon representation (left panels). Domains are colored according to Supplementary Figures S2, S3 and S4. Residues involved in LACV promoter binding are shown as green or cyan sticks and are also indicated in Supplementary Alignment File. Right panels display surface electrostatics of the 3' and 5' RNA binding sites generated by APBS within PyMol.

**Supplementary Figure S7. Potential RNA secondary structures of the 5' terminal sequences of SFTSV S, S9A, M and L segment.**

The program Mfold (2) was used to calculate potential secondary structures of the 5' RNAs of the S segment (**A**), the M segment (**B**) and the L segment (**C**). Watson-Crick base pairing is indicated by a connection between the bases.

**Supplementary Figure S8. Alignment of the 5' and 3' terminal regions of SFTSV genome segments.**

The 20 terminal nucleotides of both the 3' and the 5' genomic ends of L, M and S segments have been aligned using ClustalW (3). The figure has been created using ESPript (http://espript.ibcp.fr) (4).

**Supplementary Figure S9. Comparison of L protein polymerase activity on the different promoters of the S, M, and L segments.**

SFTSV L (D112A) protein was incubated with the conserved 5' and/or 3' terminal 20 nt (21 nt for S9A and S9U) of the S, S9A/S9U, M and L segments (Supplementary Table S1) in the presence of NTPs supplemented with [α]$^{32}$P-GTP for 60 min at 30°C. Products were separated by denaturing gel electrophoresis and visualized by autoradiography.

**Supplementary Figure S10. Additional results from biochemical *in vitro* assays.**

(**A**) The potential influence of a 5' single nucleotide C overhang on the polymerase activity of SFTSV L (D112A) was tested. The protein was incubated with the 5' vRNA either with or without an additional C and/or 3' terminal 20 nt of the M segment (Supplementary Table S1) under standard polymerase assay conditions (see Materials and Methods). Products were separated by denaturing gel electrophoresis and visualized by autoradiography. (**B**) A ssRNA oligonucleotide (Cy3-RNA) (Supplementary Table S1) 3' labelled with Cy3 (blue asterisks) was mixed in a 1:1 ratio with a ssRNA in polymerase assay buffer. The second RNA was either perfectly complementary or had missmatches ('comp' or 'miss', Supplementary Table S1) at two positions, indicated by red dots. After incubation at 30°C for 30 min the samples were mixed with an equal volume of RNA loading buffer (98% formamide, 18 mM EDTA, 0.025 mM SDS) and heated at 95°C for 3 min before they were separated on denaturing 7M Urea, 20% polyacrylamide Tris-borate-EDTA gels and 0.5-fold Tris-borate buffer at the indicated temperatures. Migration of single-stranded polyA 27mer and 40mer RNAs are shown as a size reference. (**C**) 250 nM of wild-type L protein was incubated with ~0.3 µM of radioactively labeled PolyA40 RNA substrate (Supplementary Table S1) in presence of 5 and 25 µM of the indicated Me(II) ions at 37°C for 30 min. Reactions without L protein, with EDTA or the known endonuclease-specific inhibitor DPBA added were used as negative controls. Reaction products were separated on a denaturing polyacrylamide gel and visualized by autoradiography.

**Supplementary Figure S11. Ligand plot for $m^7$GTP interaction with the CBD.**

A ligand plot for interaction of $m^7$GTP with SFTSV CBD is presented as generated by PDBsum (1). A detailed list of interactions is given in Supplementary Table S4.

**Supplementary Figure S12. Additional thermal stability data of SFTSV, RVFV and influenza virus CBDs.**

(**A**) Thermal stability of influenza virus PB2 CBD and RVFV CBD was tested in the presence or absence of different concentrations (2.5, 5.0, 7.5, and 10 mM) of $m^7$GTP, $m^7$GpppG, GTP and ATP. Melting temperatures ($T_m$) are presented as mean and standard deviations of three independent measurements (n=3). (**B**) Thermal stability of SFTSV wild type and single site mutant CBD proteins (F1703A, Y1719A, and Q1707A) is shown in the presence or absence of different concentrations (2.0, 5.0, and 10 mM) of either the cap-analogue $m^7$GTP or the unmethylated nucleotide GTP.

**Supplementary Figure S13. Cap-dependent activities of the full length L protein.**

(**A**) SFTSV L wild-type or D112A protein was incubated with 16-nt capped or uncapped primers (Supplementary Table S1) in the standard polymerase assay (see Materials and Methods). As controls the unprimed *de novo* reaction and reactions without any L protein (-L) and NTPs (-NTP) were carried out in parallel. Products were separated by denaturing gel electrophoresis and visualized by autoradiography. Primers and products are indicated. (B) Cap-dependent endonucease activity was tested by incubating 250 nM of wild-type or mutant D112A L with ~0.3 μM of radioactively labeled PolyA27 RNA substrates containing either no cap, cap0 or cap1 at their 5' termini (Supplementary Table S1) in presence of 5 μM $MnCl_2$ at 37°C for 30 min. Reactions without L protein (-L) or in the presence of EDTA were used as negative controls. Reaction products were separated on a denaturing polyacrylamide gel and visualized by autoradiography.

**Supplementary Figure S14. SAXS data.**

(**A**) The experimental scattering curve for SFTSV L is shown. (**B**) Three SAXS envelopes representing three clusters resulting from clustering analysis of a total of 40 *ab initio* models. These envelopes are referred to as envelopes 1 (blue mesh), 2 (violet mesh), and 3 (green mesh).

**Supplementary Figure S15. Additional modelling data.**

A superposition of the SAXS envelopes of clusters 2 (violet mesh) (**A**) and 3 (green mesh) (**B**), a 5Å low-pass filtered cryo-EM map A (grey surface) and the structure model (ribbon diagram, colored according to Figure 1) are presented from two different perspectives.

**Supplementary Alignment File. Alignment of *Phenuiviridae* and *Peribunyaviridae* L protein sequences.**

L protein sequences (virus abbreviations and Uniprot-IDs listed) were aligned using PROMALS3D (5) with manual adjustments. The alignment was presented by ESPript with secondary structure information given for SFTSV L protein (PDB 6Y6K) and LACV L protein (PDB 5AMQ). Regions of the L protein are labelled and marked with different colors according to Figure 1 and Supplementary Figures S2, S3 and S4. Residues of SFTSV potentially involved in 3' and 5' RNA binding are marked with dark green and dark blue frames, respectively. Residues of LACV L protein involved in 3' and 5' RNA binding are marked with light green and light blue frames, respectively. Residues of SFTSV CBD important for CBD function are marked with teal colored frames. Conserved RdRp motifs are labelled in colors according to Figure 1D. The C-terminal 500 residues vary in sequence significantly between bunyavirus families. In order to avoid unreliable results the C-terminal 500 residues of Peribunyaviridae L proteins have been removed. All numbers given refer to SFTSV L protein sequence.

#### Supplementary Table S1. Synthetic RNA used in the assays

| | Length [nt] | Name | | Sequence | |
|---|---|---|---|---|---|
| 5' RNA | 20 | 5' L | 5' HO- | ACACAGAGACGCCCAGAUGA | -OH 3' |
| | 12 | 5' L (1-12) | 5' HO- | ACACAGAGACGC | -OH 3' |
| | 10 | 5' L (1-10) | 5' HO- | ACACAGAGAC | -OH 3' |
| | 13 | 5' L (8-20) | 5' HO- | GACGCCCAGAUGA | -OH 3' |
| | 8 | 5' L (12-20) | 5' HO- | CCCAGAUGA | -OH 3' |
| | 20 | 5' M | 5' HO- | ACACAGAGACGGCCAACAAU | -OH 3' |
| | 21 | 5' M +C | 5' HO- | CACACAGAGACGGCCAACAAU | -OH 3' |
| | 20 | 5' S | 5' HO- | ACACAAAGACCCCCUUCAUU | -OH 3' |
| | 21 | 5' $S_{9A}$ | 5' HO- | ACACAAAG**A**ACCCCCUUCAUU | -OH 3' |
| 3' RNA | 20 | 3' L | 3' HO- | UGUGUUUCUGGCGGGUCUAG | -OH 5' |
| | 20 | 3' M | 3' HO- | UGUGUUUCUGGCCGGUUGUG | -OH 5' |
| | 20 | 3' S | 3' HO- | UGUGUUUCUGGGGGUUUUUU | -OH 5' |
| | 21 | 3' $S_{9U}$ | 3' HO- | UGUGUUUC**U**UGGGGGUUUUUU | -OH 5' |
| Primer | 16 | Primer 1 | 5' PPP- | AAACGCAACAACAACA | -OH 3' |
| | 16 | Primer 2 | 5' HO- | AAACGCAACAACAACA | -OH 3' |
| | | Primer 3 | 5' HO- | ACACAAA | -OH 3' |
| PolyA | 27 | PA27-P3 | 5' PPP- | $(A)_n$ (n=27) | -OH 3' |
| | 27 | PA27-OH | 5'HO- | $(A)_n$ (n=27) | -OH 3' |
| | 40 | PA40-OH | 5'HO- | $(A)_n$ (n=40) | -OH 3' |
| dsRNA | 20 | cy3-RNA | 5'HO- | GCGCACCGGGGAUCCUAGGC | -cy3 |
| | 19 | miss | 3' HO- | GCGUG**U**C**A**CCUAGGAUCCG | -OH 5' |
| | 19 | comp | 3' HO- | GCGUGGCCCCUAGGAUCCG | -OH 5' |

The table lists the RNA oligonucleotides that were used in the described assays. The specific sequence, length in nucleotides and the identifier used to label the RNA in the experimental descriptions are given. RNAs were synthesized by Biomers.

**Supplementary Table S2. CryoEM data collection, refinement and validation statistics.**

| **DATA COLLECTION AND PROCESSING** | Map A |
|---|---|
| Magnification | 105,000 |
| Voltage (kV) | 300 |
| Electron exposure (e-/Å²) | 24.8 |
| Defocus range (μm) | 1-3 |
| Pixel size (Å) | 0.87 |
| Symmetry imposed | C1 |
| Initial particle images (no.) | 779,935 |
| Final particle images (no.) | 223,063 |
| Map resolution (Å) | 3.8 |
| FSC threshold | 0.143 |
| Map resolution range | 3.3 – 4.8 |
| **REFINEMENT** | |
| Initial model used | 5AMR |
| Model resolution (Å) | 3.7 |
| FSC threshold | 0.5 |
| Map sharpening B factor (Å²) | -224 |
| Model versus map cross-correlation | 0.88 |
| Model composition: | |
| Nonhydrogen atoms | 10351 |
| Protein residues | 1299 |
| Ligand | 1 |
| R.m.s deviations: | |
| Bond lengths (Å) | 0.012 |
| Bond angles (°) | 0.982 |
| Validation: | |
| MolProbity score | 2.45 |
| Clashscore | 10.98 |
| Poor rotamers (%) | 2.36 |
| Ramachandran plot: | |
| Favored (%) | 87.37 |
| Allowed (%) | 12.47 |
| Disallowed (%) | 0.16 |
| PDB code | 6Y6K |
| EMDB code | EMD-10706 |

**Supplementary Table S3. Crystallographic data and refinement statistics.**

| | **SFTSV CBD** |
|---|---|
| **Data collection** | |
| Synchrotron beamline | PETRA P14 |
| Wavelength (Å) | 0.9763 |
| Resolution (Å) | 50 - 1.48 (1.398 - 1.35) |
| Space group | $P2_12_12_1$ |
| Cell dimensions | |
| a, b, c (Å) | 38.974, 44.715 63.515 |
| α, β, γ (°) | 90, 90, 90 |
| Total reflections | 321482 (25283) |
| Unique reflections | 24447 (2183) |
| Multiplicity | 13.2 (11.6) |
| Completeness (%) | 0.976 (0.888) |
| Mean I/sigma(I) | 41.6 (6.32) |
| Wilson B-factor | 16.17 |
| R-merge | 0.03094 (0.3684) |
| R-meas | 0.03223 (0.3857) |
| **Refinement** | |
| Resolution (Å) | 33.22-1.35 (1.398-1.35) |
| Reflections used in refinement | 24438 (2182) |
| Reflections used for R-free | 1188 (118) |
| R-work | 0.1656 (0.2209) |
| R-free | 0.1764 (0.2544) |
| No. of atoms | |
| protein | 917 |
| ligand/ ion | 34 |
| Water molecules | 189 |
| Average B-factors ($Å^2$) | |
| protein | 18.99 |
| ligand | 25.11 |
| solvent | 34.04 |
| R.m.s deviations | |
| bond lengths (Å) | 0.008 |
| bond angles (°) | 0.95 |
| Ramachandran (%) | |
| favored | 100 |
| allowed | 0 |
| outliers | 0 |
| Rotamer outliers (%) | 0 |
| **PDB code** | **6XYA** |

#### Supplementary Table S4. List of interactions of SFTSV CBD with m⁷GTP generated by PDBsum.

**Hydrogen bonds**

|  | <---- | ATOM 1 |  |  | -----> |  | <--- | ATOM 2 |  |  | -----> |  |
|---|---|---|---|---|---|---|---|---|---|---|---|---|
| | Atom | Atom | Res | Res | | | Atom | Atom | Res | Res | | |
| | no. | name | name | no. | Chain | | no. | name | name | no. | Chain | Distance |
| 1 | 121 | N | GLN | 1707 | B | --> | 1871 | O6 | MGP | 1 | A | 3.03 |
| 2 | 128 | OE1 | GLN | 1707 | B | <-- | 1853 | N1 | MGP | 1 | A | 2.62 |
| 3 | 128 | OE1 | GLN | 1707 | B | <-- | 1854 | N2 | MGP | 1 | A | 2.97 |
| 4 | 1175 | O | ASP | 1771 | B | <-- | 1861 | O2' | MGP | 1 | A | 2.9 |
| 5 | 1187 | O | LEU | 1772 | B | <-- | 1854 | N2 | MGP | 1 | A | 3.01 |

**Non-bonded contacts**

|  | <---- | ATOM 1 |  |  | -----> |  | <--- | ATOM 2 |  |  | -----> |  |
|---|---|---|---|---|---|---|---|---|---|---|---|---|
| | Atom | Atom | Res | Res | | | Atom | Atom | Res | Res | | |
| | no. | name | name | no. | Chain | | no. | name | name | no. | Chain | Distance |
| 1 | 60 | CD1 | PHE | 1703 | B | --- | 1871 | O6 | MGP | 1 | A | 3.84 |
| 2 | 61 | CD2 | PHE | 1703 | B | --- | 1852 | CM7 | MGP | 1 | A | 3.88 |
| 3 | 61 | CD2 | PHE | 1703 | B | --- | 1858 | O1A | MGP | 1 | A | 3.65 |
| 4 | 62 | CE1 | PHE | 1703 | B | --- | 1850 | C6 | MGP | 1 | A | 3.67 |
| 5 | 62 | CE1 | PHE | 1703 | B | --- | 1853 | N1 | MGP | 1 | A | 3.8 |
| 6 | 62 | CE1 | PHE | 1703 | B | --- | 1871 | O6 | MGP | 1 | A | 3.67 |
| 7 | 63 | CE2 | PHE | 1703 | B | --- | 1848 | C5 | MGP | 1 | A | 3.52 |
| 8 | 63 | CE2 | PHE | 1703 | B | --- | 1850 | C6 | MGP | 1 | A | 3.78 |
| 9 | 63 | CE2 | PHE | 1703 | B | --- | 1856 | N7 | MGP | 1 | A | 3.66 |
| 10 | 64 | CZ | PHE | 1703 | B | --- | 1848 | C5 | MGP | 1 | A | 3.61 |
| 11 | 64 | CZ | PHE | 1703 | B | --- | 1850 | C6 | MGP | 1 | A | 3.43 |
| 12 | 64 | CZ | PHE | 1703 | B | --- | 1853 | N1 | MGP | 1 | A | 3.61 |
| 13 | 64 | CZ | PHE | 1703 | B | --- | 1871 | O6 | MGP | 1 | A | 3.8 |
| 14 | 121 | N | GLN | 1707 | B | --- | 1871 | O6 | MGP | 1 | A | 3.03 |
| 15 | 122 | CA | GLN | 1707 | B | --- | 1871 | O6 | MGP | 1 | A | 3.79 |
| 16 | 125 | CB | GLN | 1707 | B | --- | 1871 | O6 | MGP | 1 | A | 3.53 |
| 17 | 126 | CG | GLN | 1707 | B | --- | 1871 | O6 | MGP | 1 | A | 3.86 |
| 18 | 127 | CD | GLN | 1707 | B | --- | 1853 | N1 | MGP | 1 | A | 3.61 |
| 19 | 128 | OE1 | GLN | 1707 | B | --- | 1843 | C2 | MGP | 1 | A | 3.23 |
| 20 | 128 | OE1 | GLN | 1707 | B | --- | 1850 | C6 | MGP | 1 | A | 3.72 |
| 21 | 128 | OE1 | GLN | 1707 | B | --- | 1853 | N1 | MGP | 1 | A | 2.62 |
| 22 | 128 | OE1 | GLN | 1707 | B | --- | 1854 | N2 | MGP | 1 | A | 2.97 |
| 23 | 296 | CB | TYR | 1719 | B | --- | 1843 | C2 | MGP | 1 | A | 3.75 |
| 24 | 296 | CB | TYR | 1719 | B | --- | 1853 | N1 | MGP | 1 | A | 3.58 |
| 25 | 297 | CG | TYR | 1719 | B | --- | 1843 | C2 | MGP | 1 | A | 3.62 |
| 26 | 297 | CG | TYR | 1719 | B | --- | 1846 | C4 | MGP | 1 | A | 3.81 |
| 27 | 297 | CG | TYR | 1719 | B | --- | 1848 | C5 | MGP | 1 | A | 3.79 |
| 28 | 297 | CG | TYR | 1719 | B | --- | 1850 | C6 | MGP | 1 | A | 3.73 |
| 29 | 297 | CG | TYR | 1719 | B | --- | 1853 | N1 | MGP | 1 | A | 3.63 |
| 30 | 297 | CG | TYR | 1719 | B | --- | 1855 | N3 | MGP | 1 | A | 3.75 |
| 31 | 298 | CD1 | TYR | 1719 | B | --- | 1846 | C4 | MGP | 1 | A | 3.81 |
| 32 | 298 | CD1 | TYR | 1719 | B | --- | 1848 | C5 | MGP | 1 | A | 3.44 |

| | | | | | | | | | | | | |
|---|---|---|---|---|---|---|---|---|---|---|---|---|
| 33 | 298 | CD1 | TYR | 1719 | B | --- | 1850 | C6 | MGP | 1 | A | 3.53 |
| 34 | 298 | CD1 | TYR | 1719 | B | --- | 1856 | N7 | MGP | 1 | A | 3.8 |
| 35 | 299 | CD2 | TYR | 1719 | B | --- | 1843 | C2 | MGP | 1 | A | 3.69 |
| 36 | 299 | CD2 | TYR | 1719 | B | --- | 1846 | C4 | MGP | 1 | A | 3.64 |
| 37 | 299 | CD2 | TYR | 1719 | B | --- | 1855 | N3 | MGP | 1 | A | 3.41 |
| 38 | 300 | CE1 | TYR | 1719 | B | --- | 1846 | C4 | MGP | 1 | A | 3.67 |
| 39 | 300 | CE1 | TYR | 1719 | B | --- | 1848 | C5 | MGP | 1 | A | 3.45 |
| 40 | 300 | CE1 | TYR | 1719 | B | --- | 1851 | C8 | MGP | 1 | A | 3.56 |
| 41 | 300 | CE1 | TYR | 1719 | B | --- | 1856 | N7 | MGP | 1 | A | 3.4 |
| 42 | 300 | CE1 | TYR | 1719 | B | --- | 1857 | N9 | MGP | 1 | A | 3.78 |
| 43 | 301 | CE2 | TYR | 1719 | B | --- | 1842 | C1' | MGP | 1 | A | 3.65 |
| 44 | 301 | CE2 | TYR | 1719 | B | --- | 1846 | C4 | MGP | 1 | A | 3.51 |
| 45 | 301 | CE2 | TYR | 1719 | B | --- | 1855 | N3 | MGP | 1 | A | 3.66 |
| 46 | 301 | CE2 | TYR | 1719 | B | --- | 1857 | N9 | MGP | 1 | A | 3.62 |
| 47 | 302 | CZ | TYR | 1719 | B | --- | 1842 | C1' | MGP | 1 | A | 3.75 |
| 48 | 302 | CZ | TYR | 1719 | B | --- | 1846 | C4 | MGP | 1 | A | 3.52 |
| 49 | 302 | CZ | TYR | 1719 | B | --- | 1848 | C5 | MGP | 1 | A | 3.81 |
| 50 | 302 | CZ | TYR | 1719 | B | --- | 1851 | C8 | MGP | 1 | A | 3.5 |
| 51 | 302 | CZ | TYR | 1719 | B | --- | 1856 | N7 | MGP | 1 | A | 3.8 |
| 52 | 302 | CZ | TYR | 1719 | B | --- | 1857 | N9 | MGP | 1 | A | 3.36 |
| 53 | 303 | OH | TYR | 1719 | B | --- | 1842 | C1' | MGP | 1 | A | 3.66 |
| 54 | 303 | OH | TYR | 1719 | B | --- | 1851 | C8 | MGP | 1 | A | 3.49 |
| 55 | 303 | OH | TYR | 1719 | B | --- | 1857 | N9 | MGP | 1 | A | 3.49 |
| 56 | 303 | OH | TYR | 1719 | B | --- | 1869 | O4' | MGP | 1 | A | 3.5 |
| 57 | 1175 | O | ASP | 1771 | B | --- | 1844 | C2' | MGP | 1 | A | 3.77 |
| 58 | 1175 | O | ASP | 1771 | B | --- | 1861 | O2' | MGP | 1 | A | 2.9 |
| 59 | 1175 | O | ASP | 1771 | B | --- | 1865 | O3' | MGP | 1 | A | 3.79 |
| 60 | 1187 | O | LEU | 1772 | B | --- | 1843 | C2 | MGP | 1 | A | 3.88 |
| 61 | 1187 | O | LEU | 1772 | B | --- | 1854 | N2 | MGP | 1 | A | 3.01 |
| 62 | 1187 | O | LEU | 1772 | B | --- | 1855 | N3 | MGP | 1 | A | 3.81 |
| 63 | 1191 | CD2 | LEU | 1772 | B | --- | 1844 | C2' | MGP | 1 | A | 3.54 |
| 64 | 1191 | CD2 | LEU | 1772 | B | --- | 1845 | C3' | MGP | 1 | A | 3.63 |
| 65 | 1191 | CD2 | LEU | 1772 | B | --- | 1865 | O3' | MGP | 1 | A | 3.54 |

Number of hydrogen bonds: 5

Number of non-bonded contacts: 65

Supplementary Figure S1

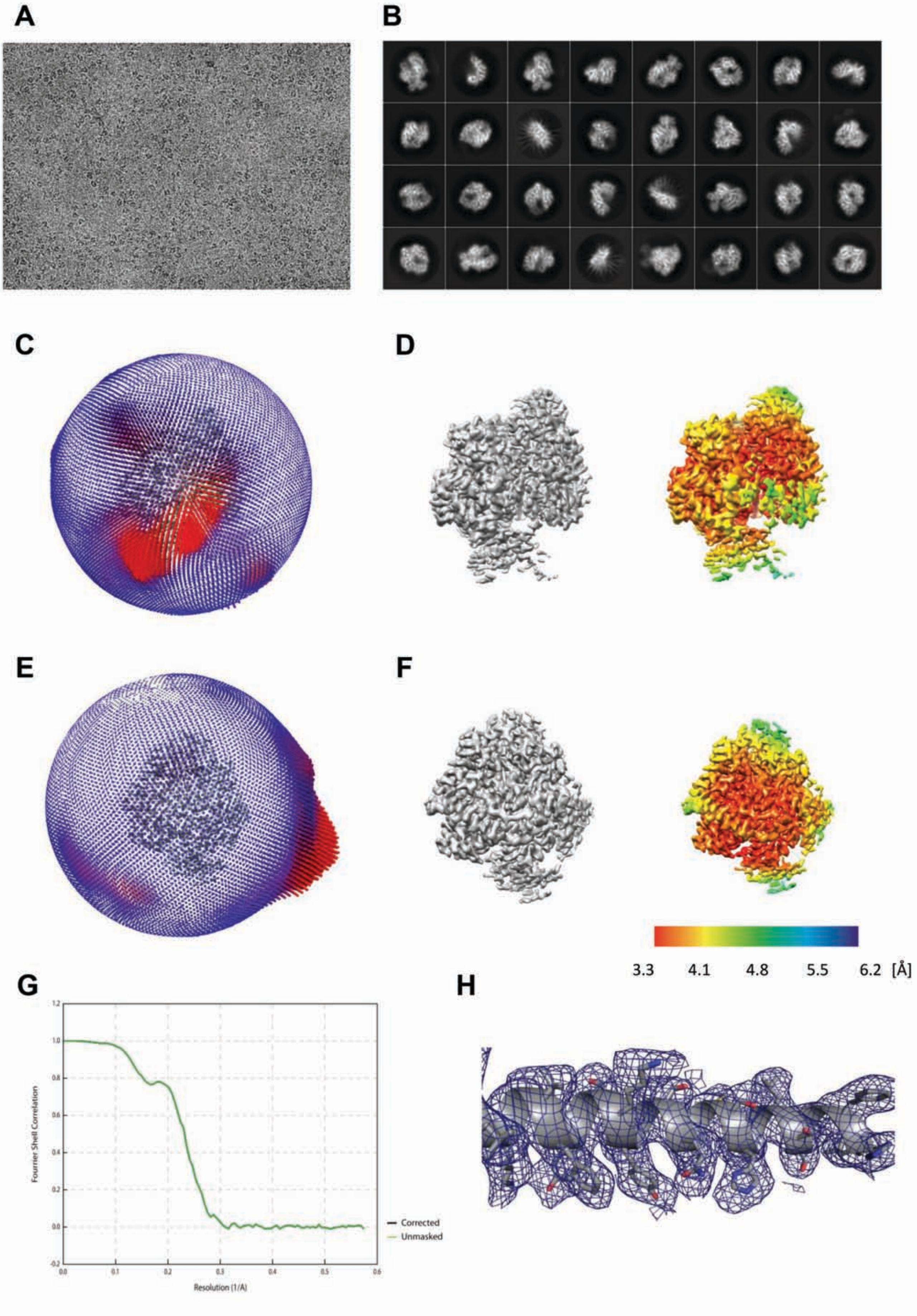

#### Supplementary Figure S2

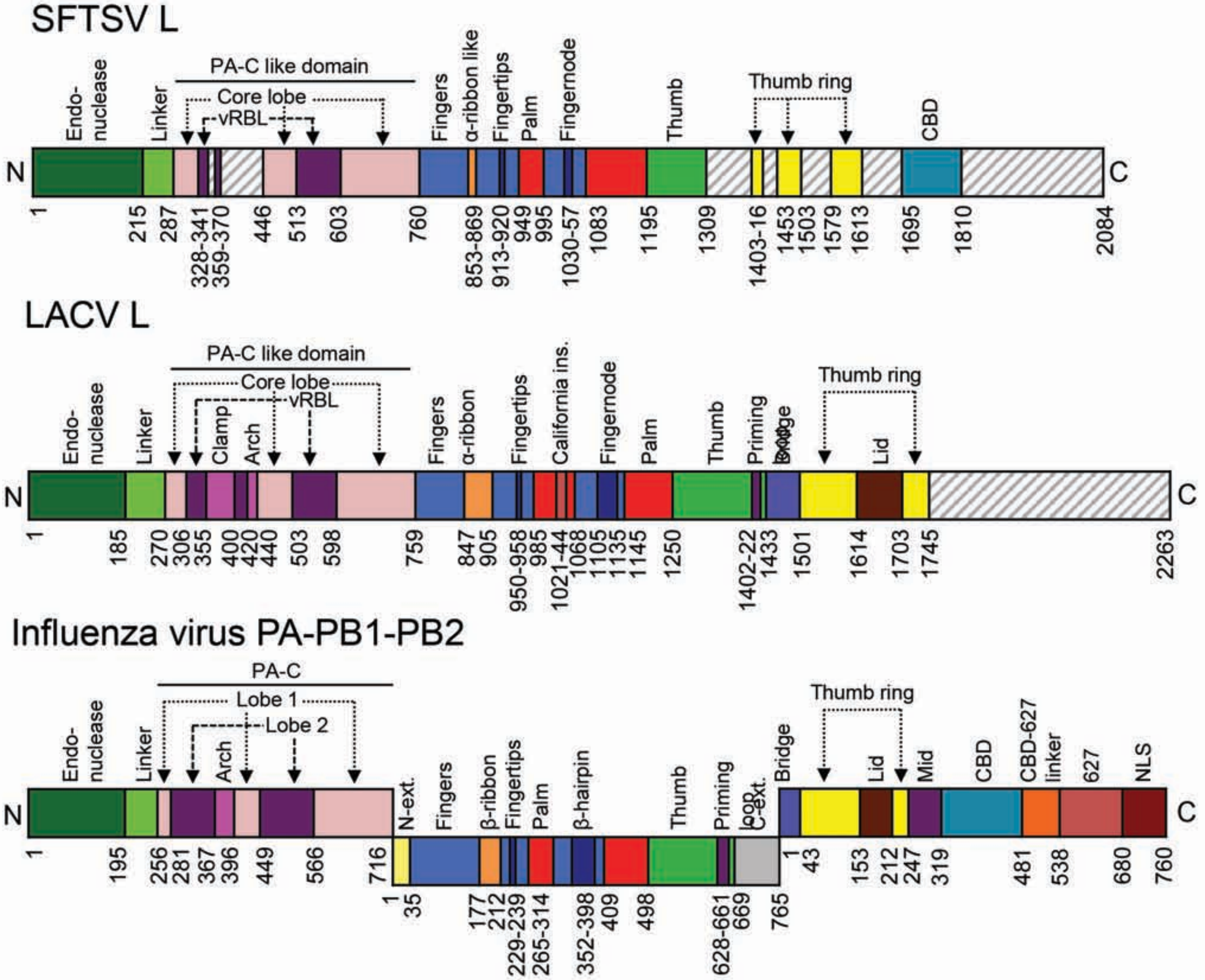

Supplementary Figure S3

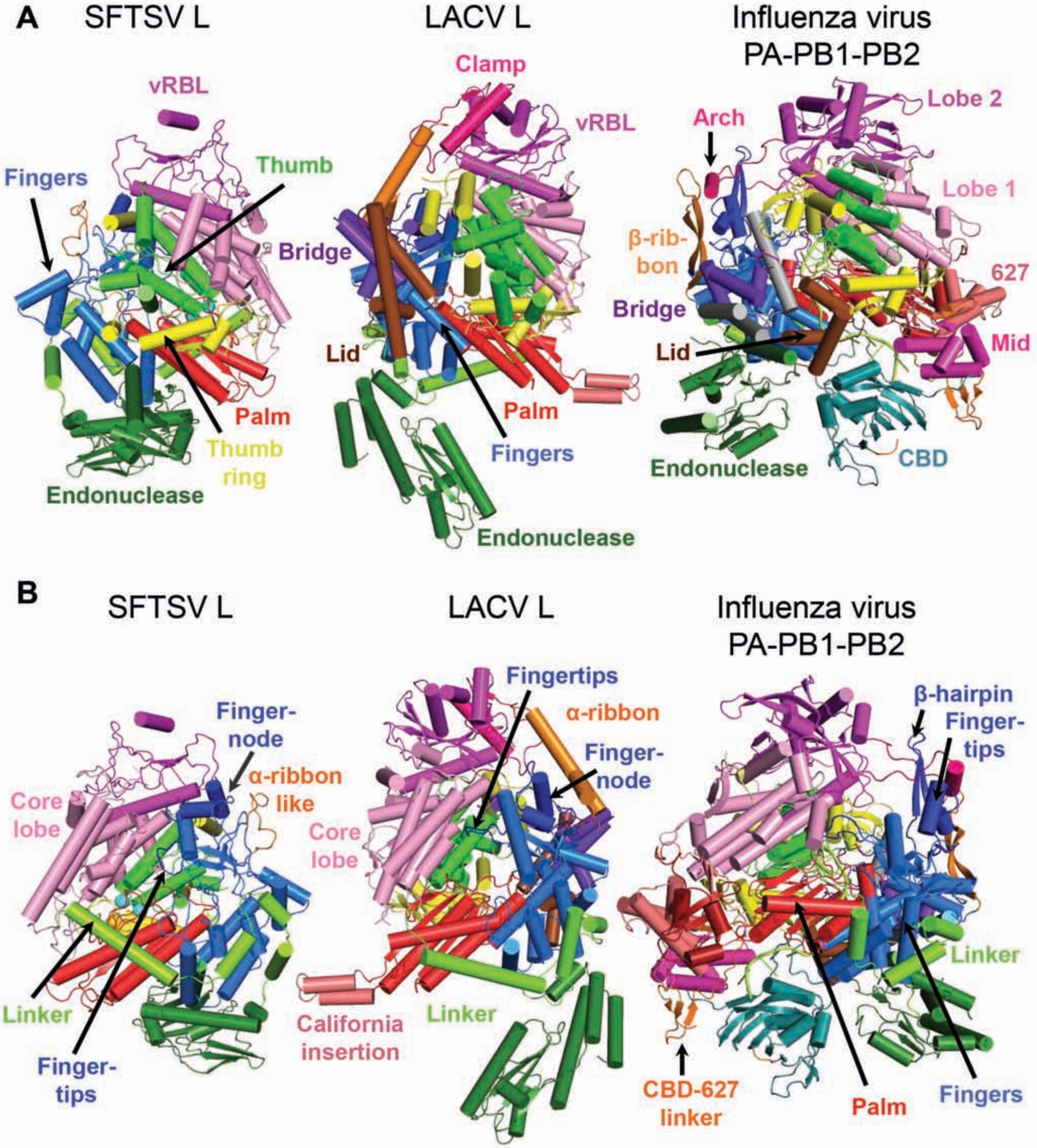

Supplementary Figure S4

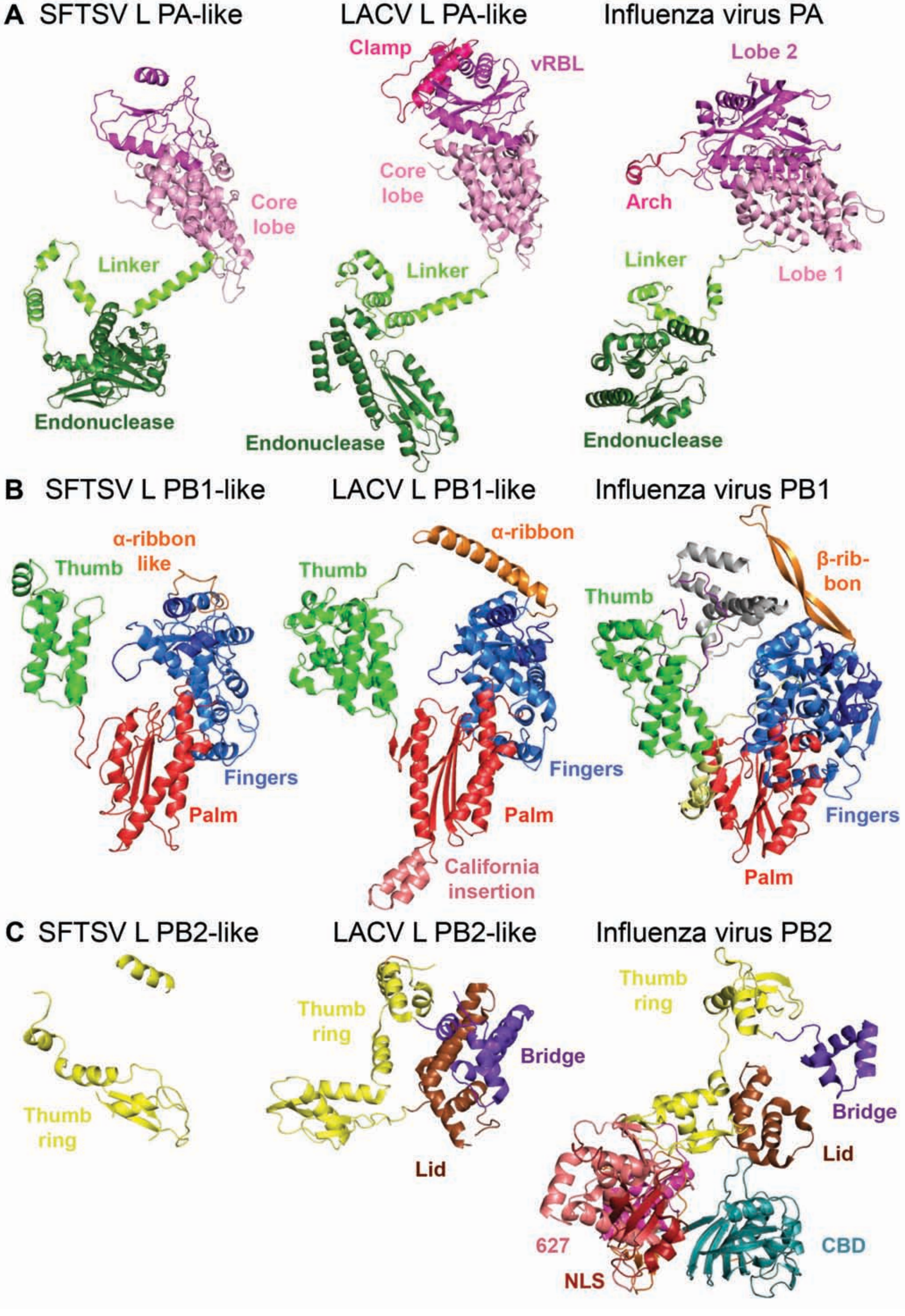

Supplementary Figure S5

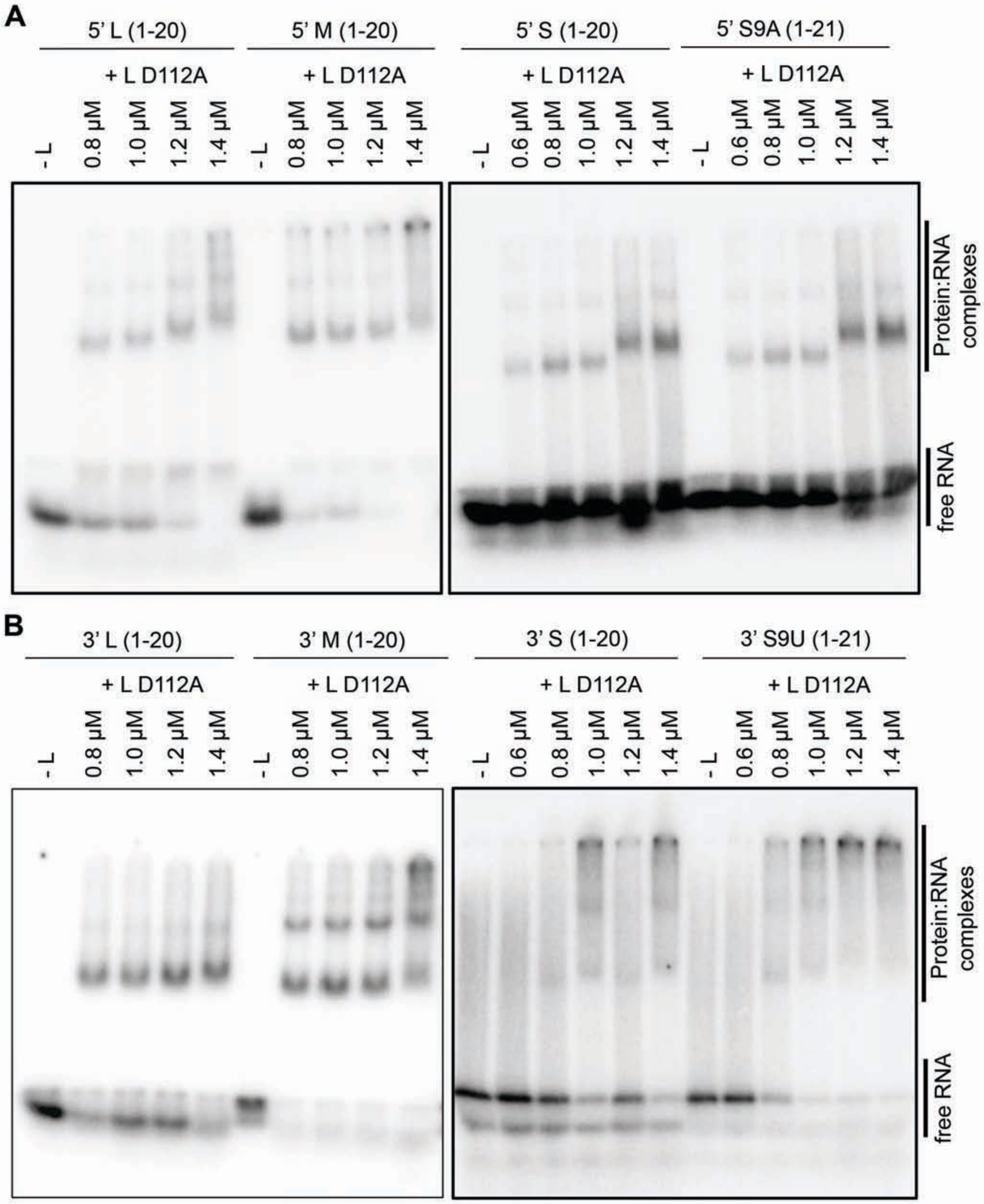

Supplementary Figure S6

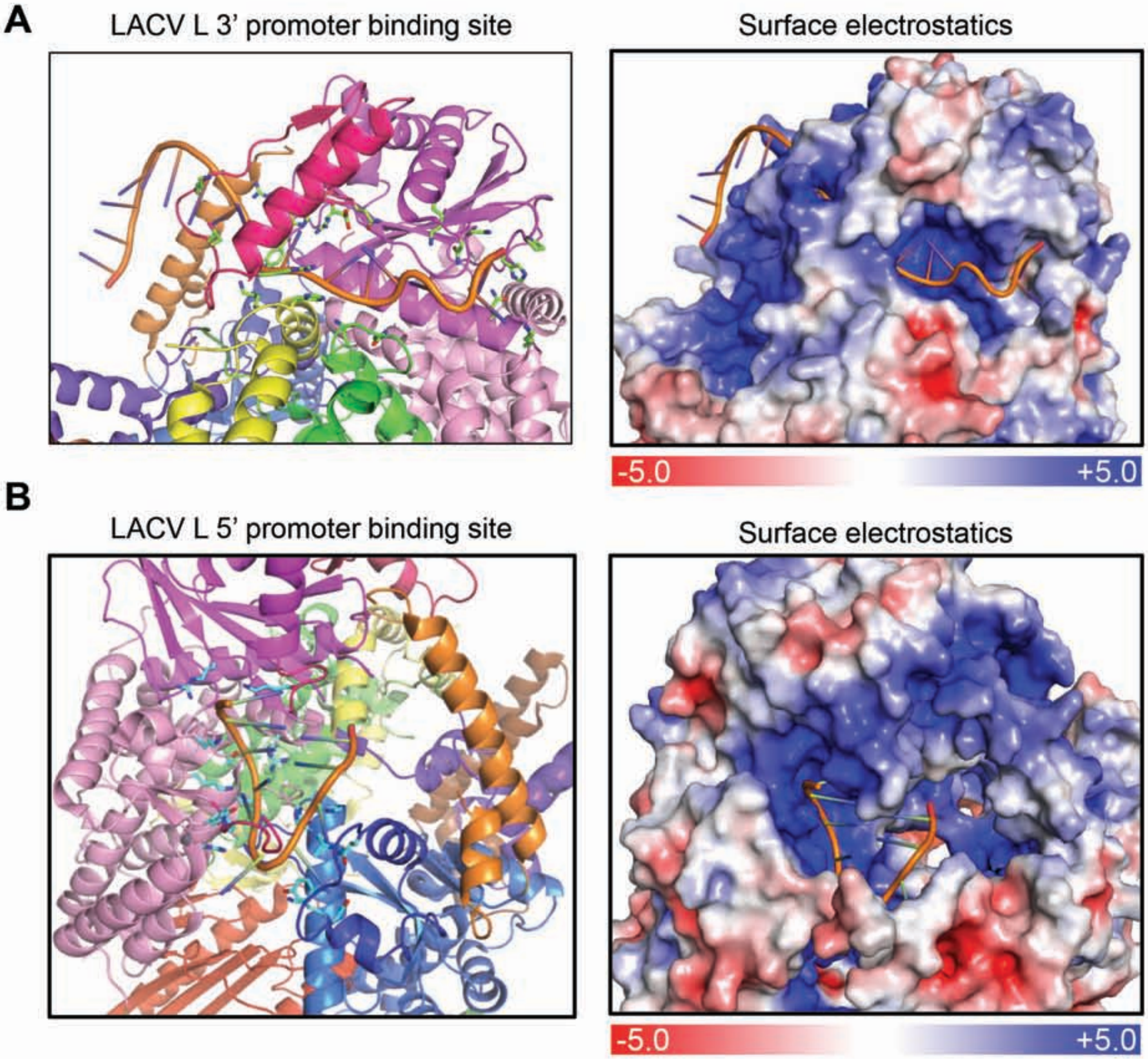

### Supplementary Figure S7

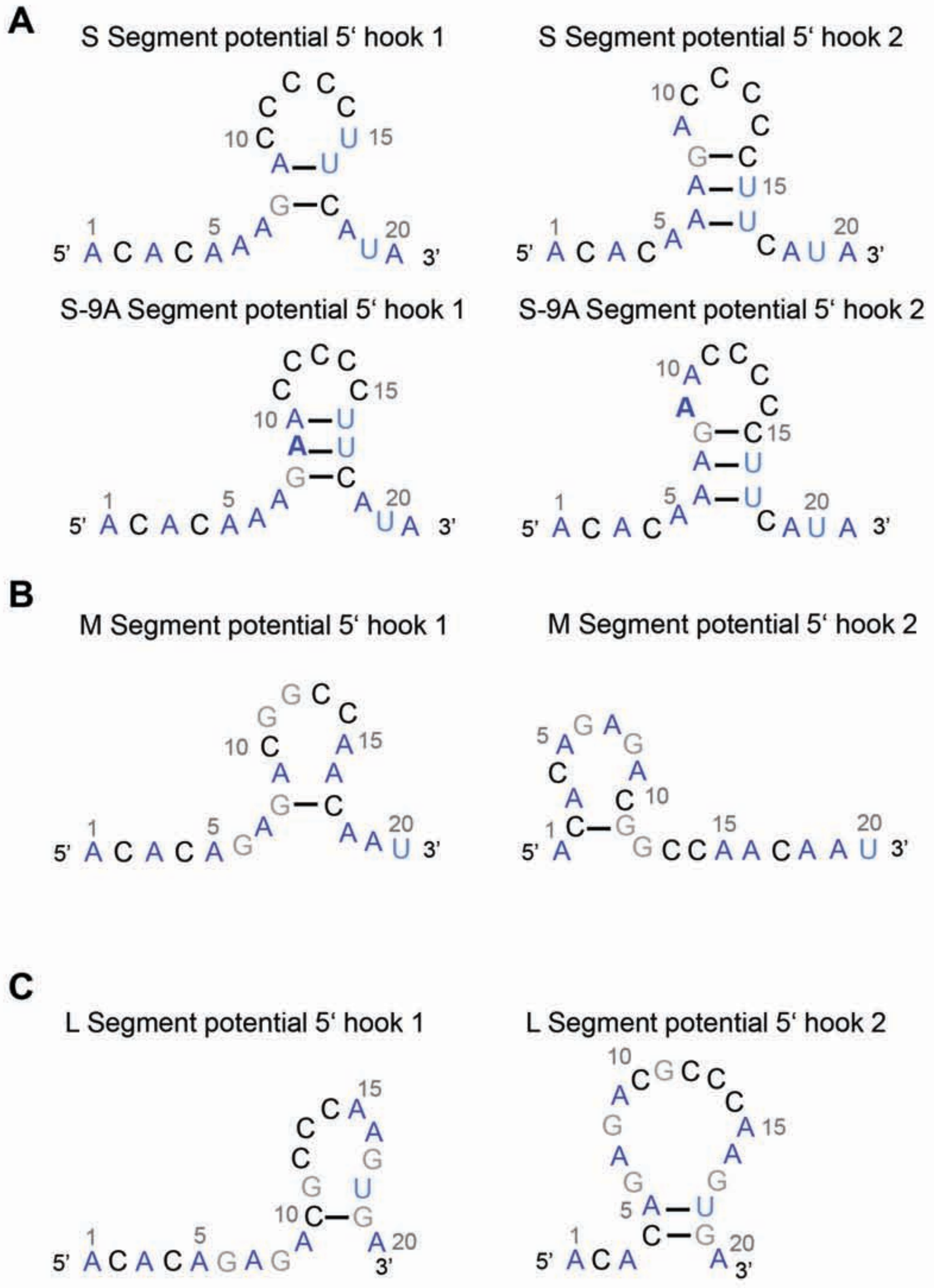

### Supplementary Figure S8

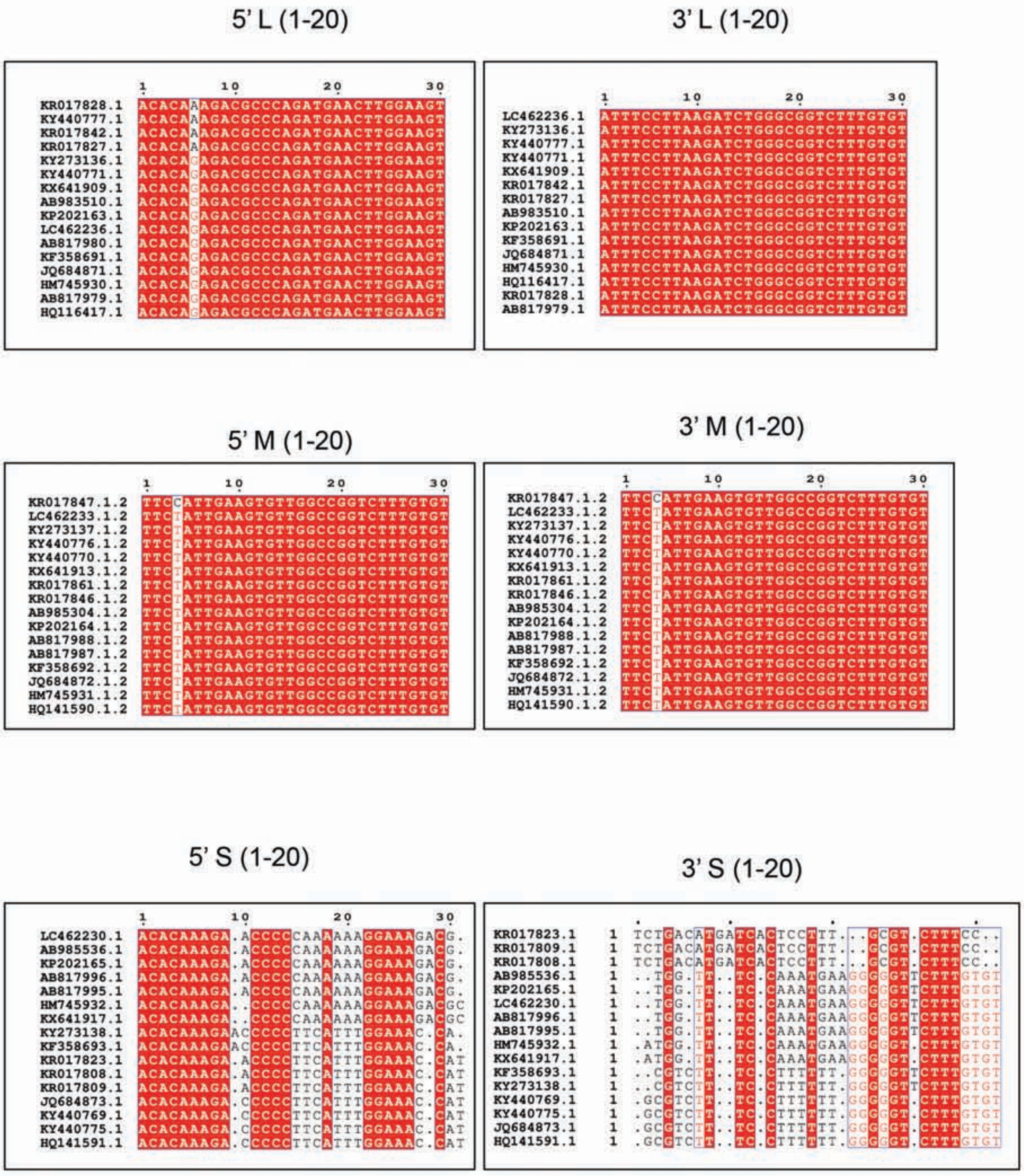

Supplementary Figure S9:

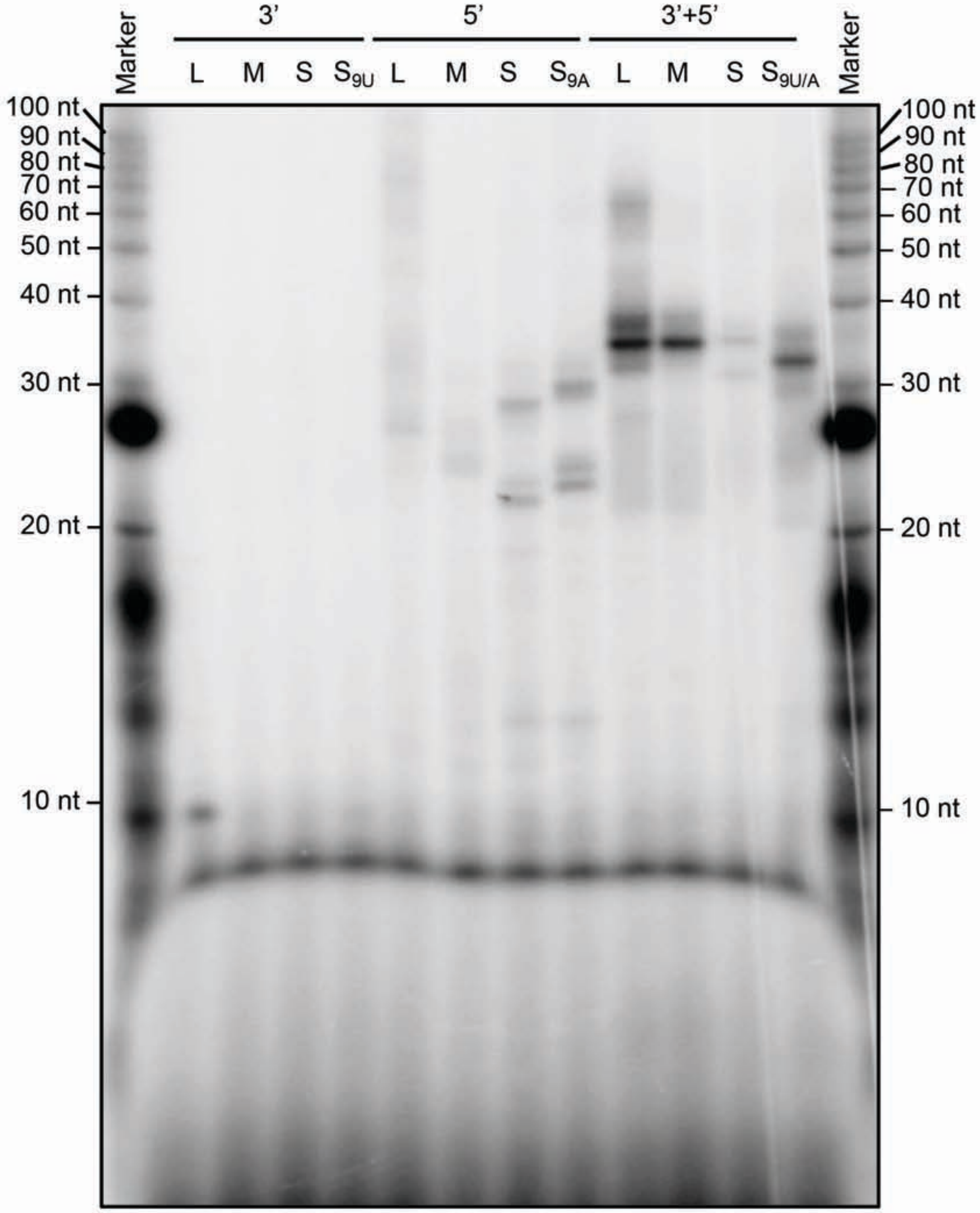

Supplementary Figure S10

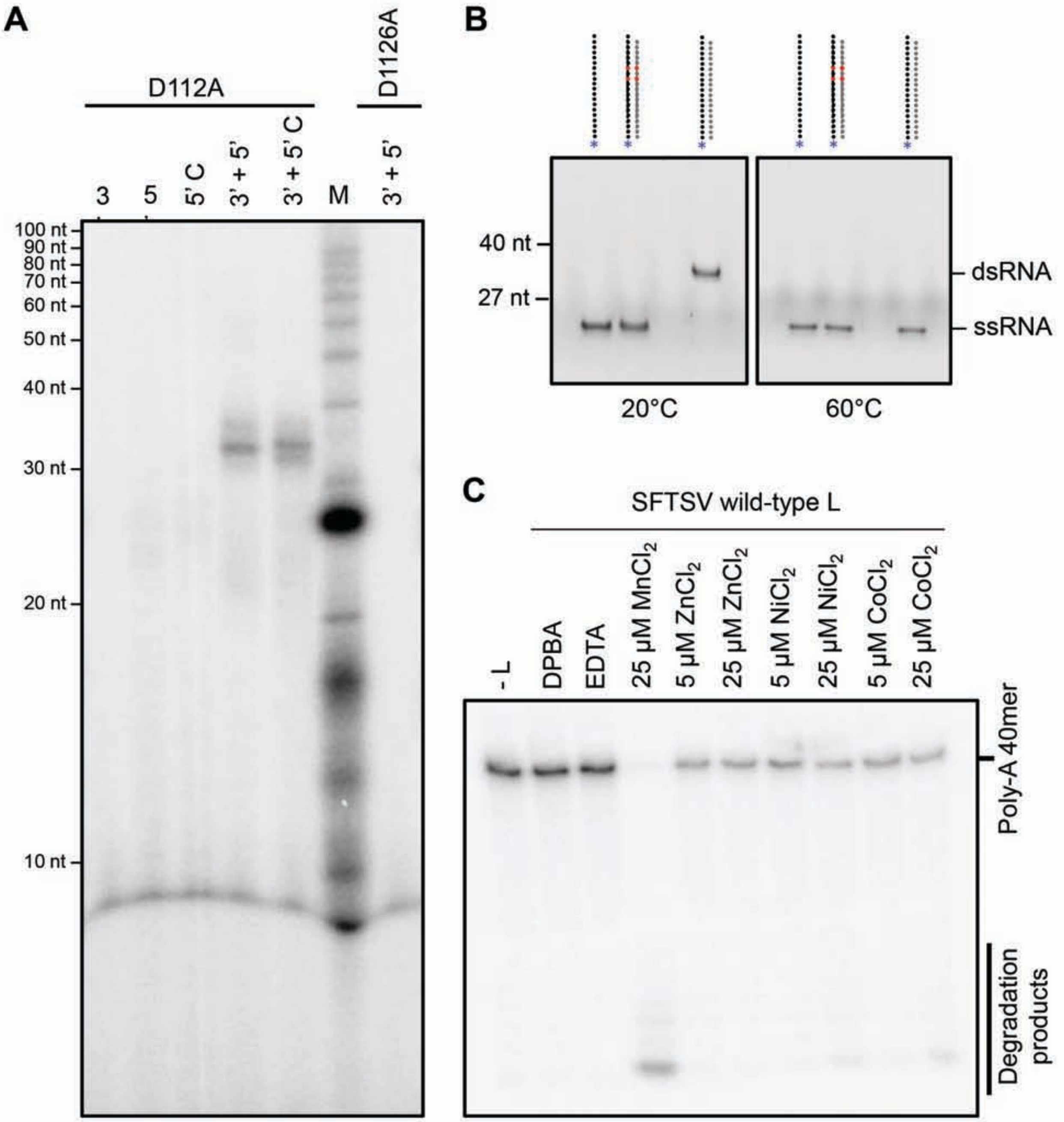

Supplementary Figure S11

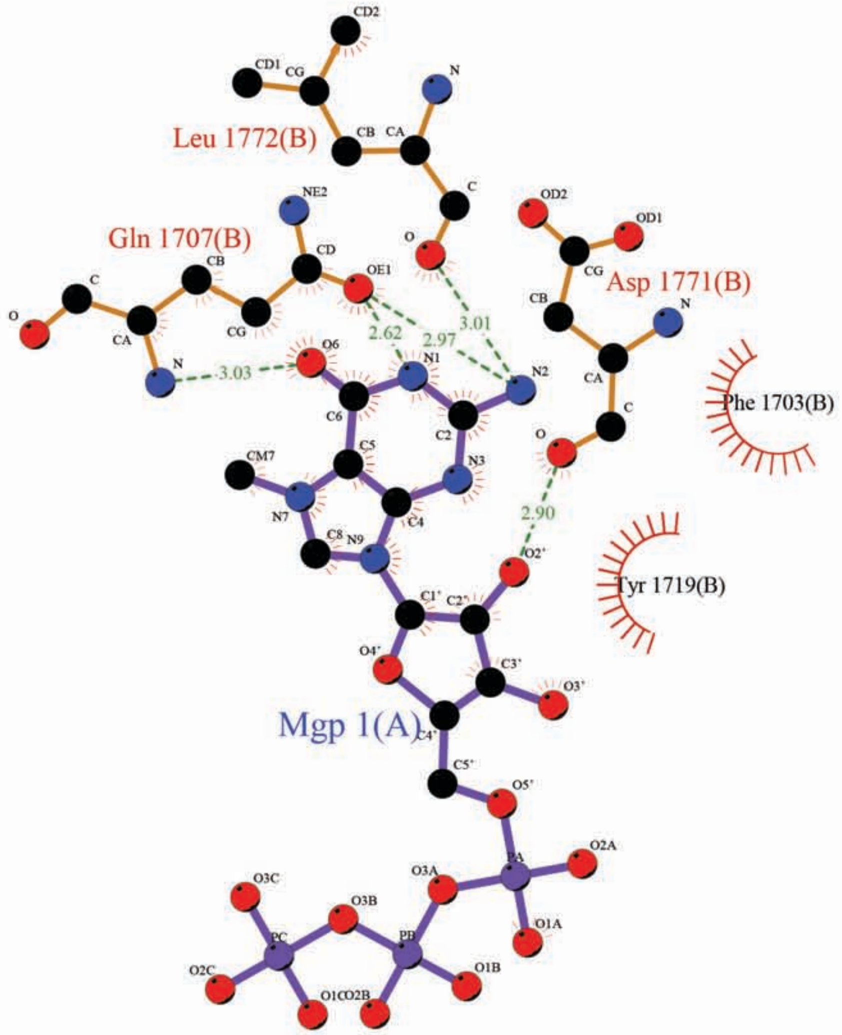

### Supplementary Figure S12

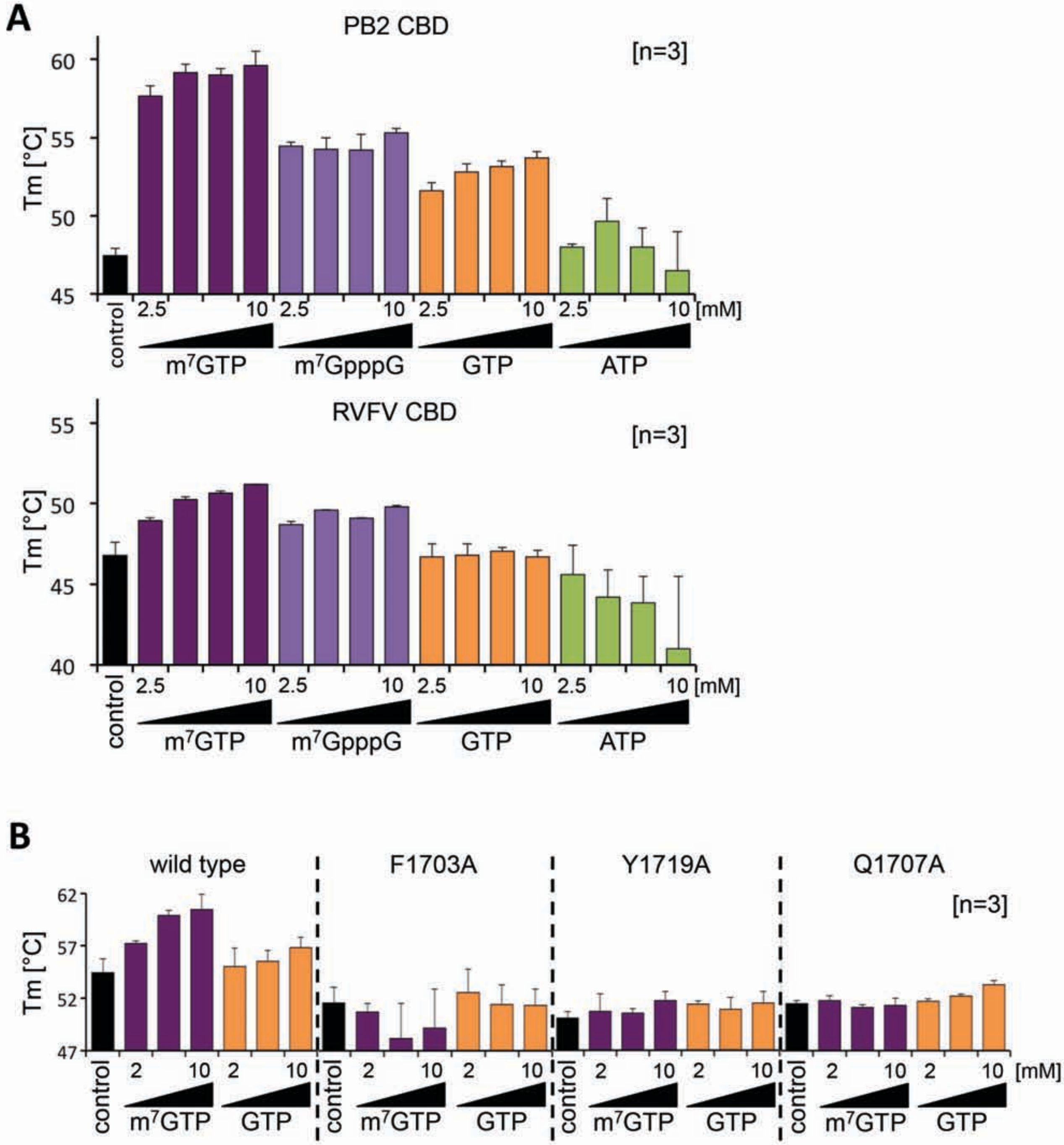

Supplementary Figure S13

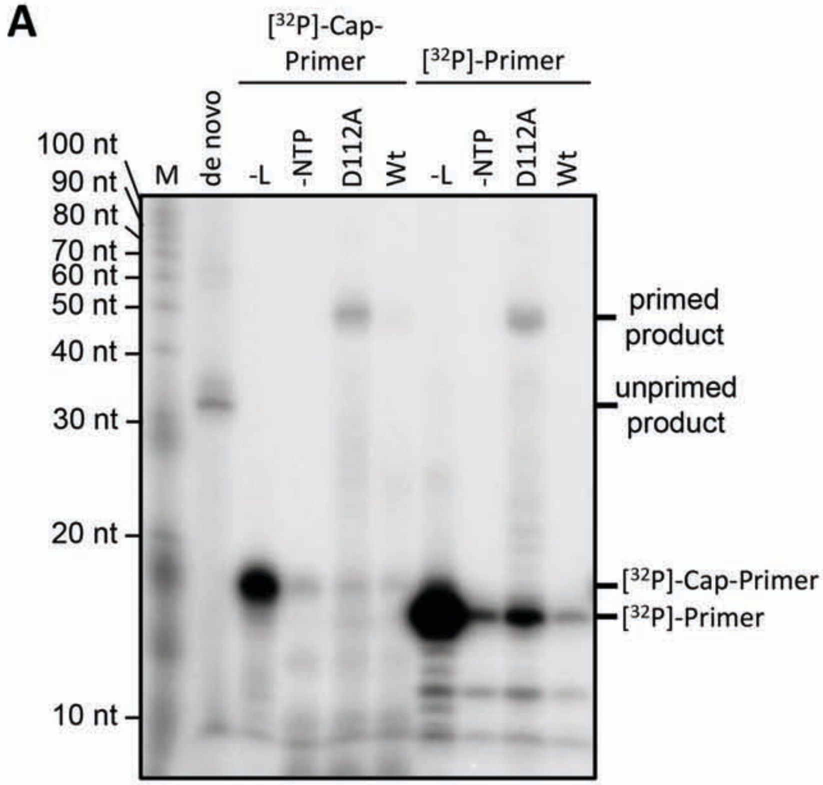


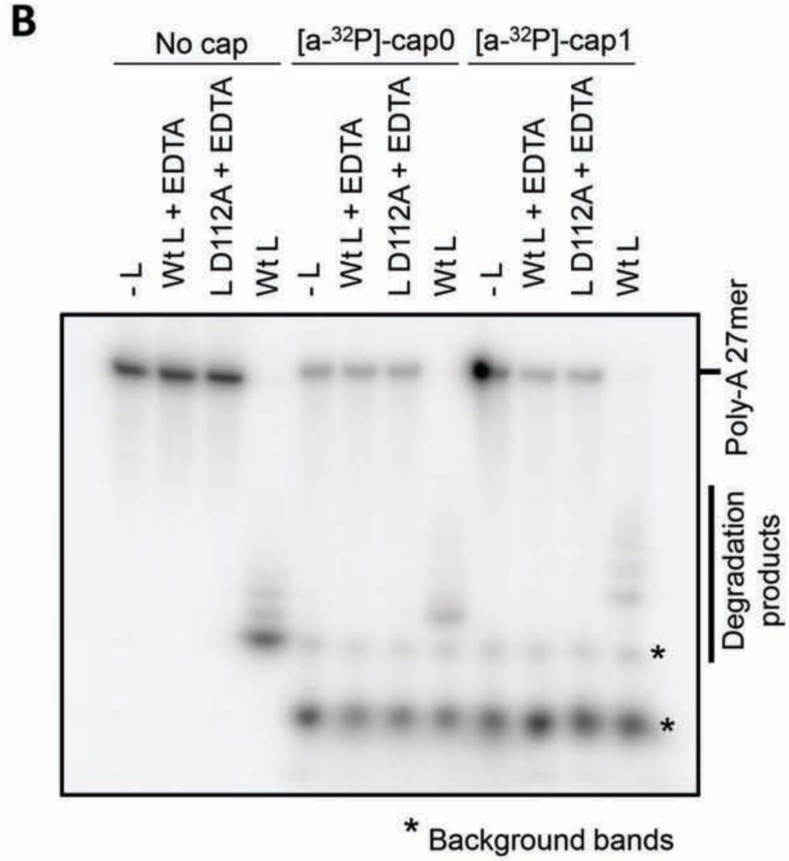

Supplementary Figure S14

A

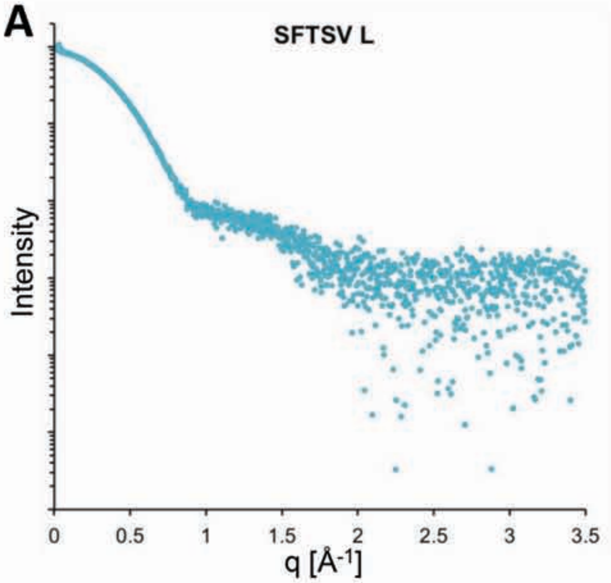


B

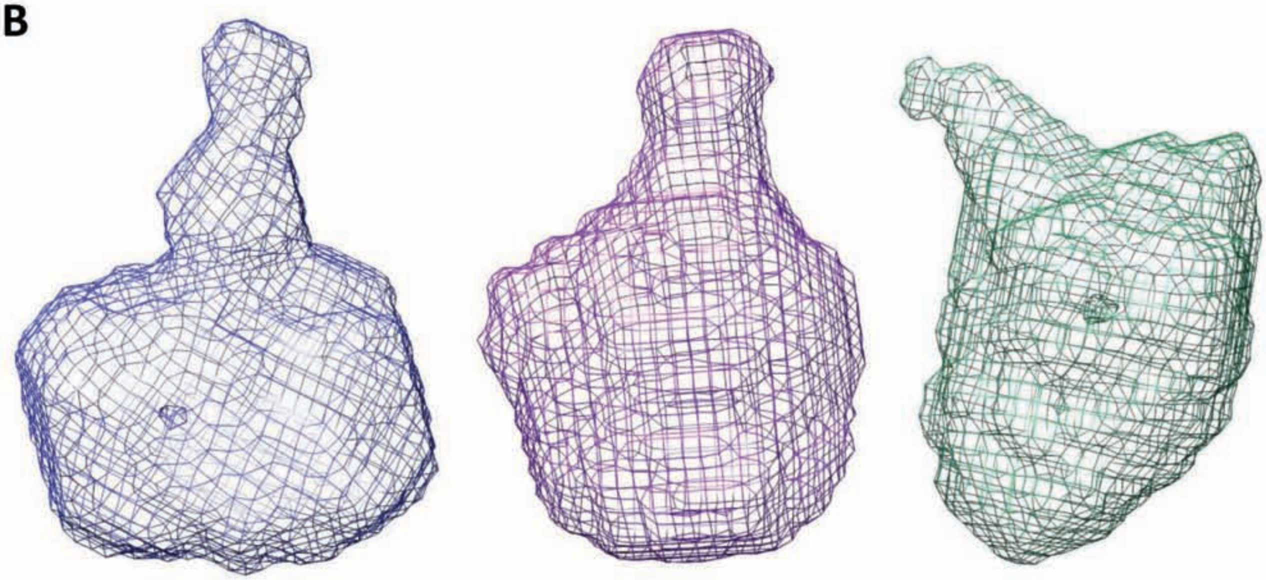

Supplementary Figure S15

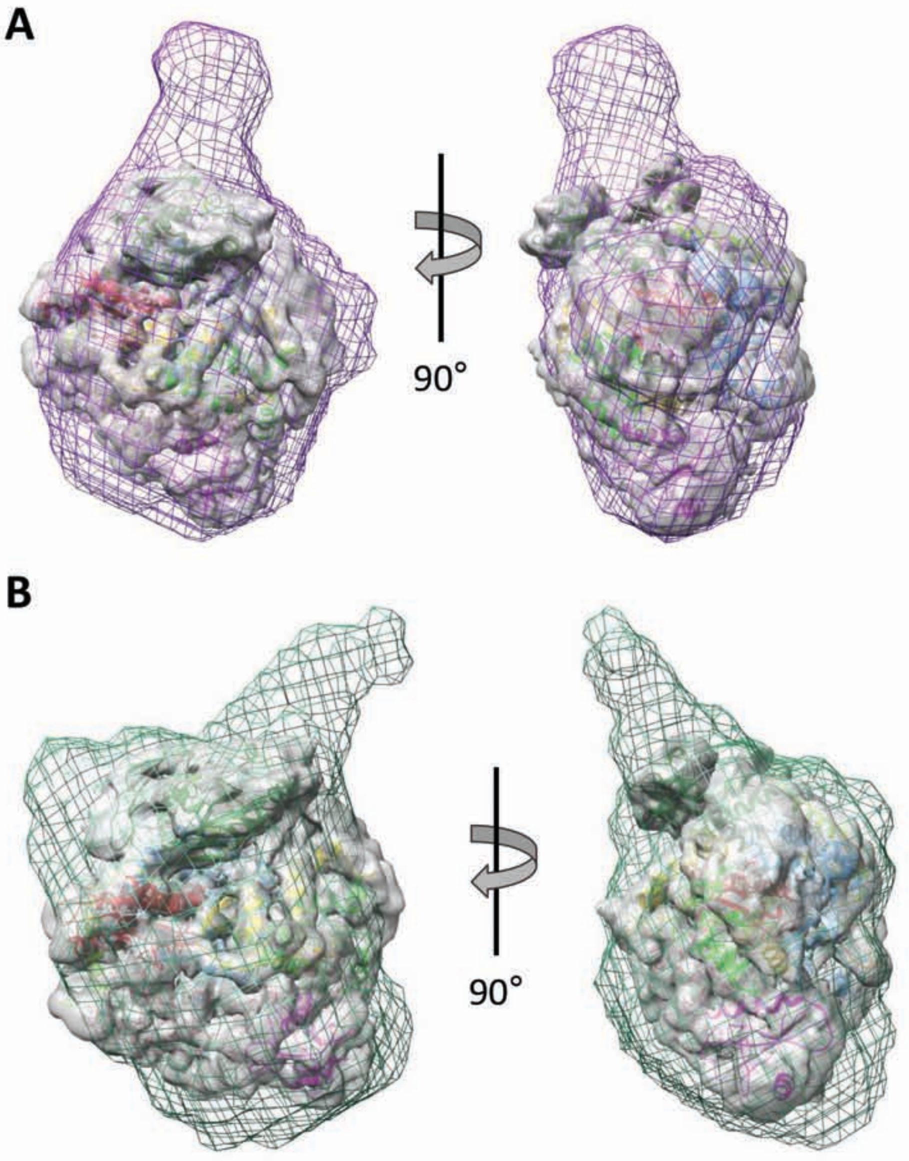
