## Supplementary Alignment File for "Structural and functional characterization of the Severe fever with thrombocytopenia syndrome virus L protein"

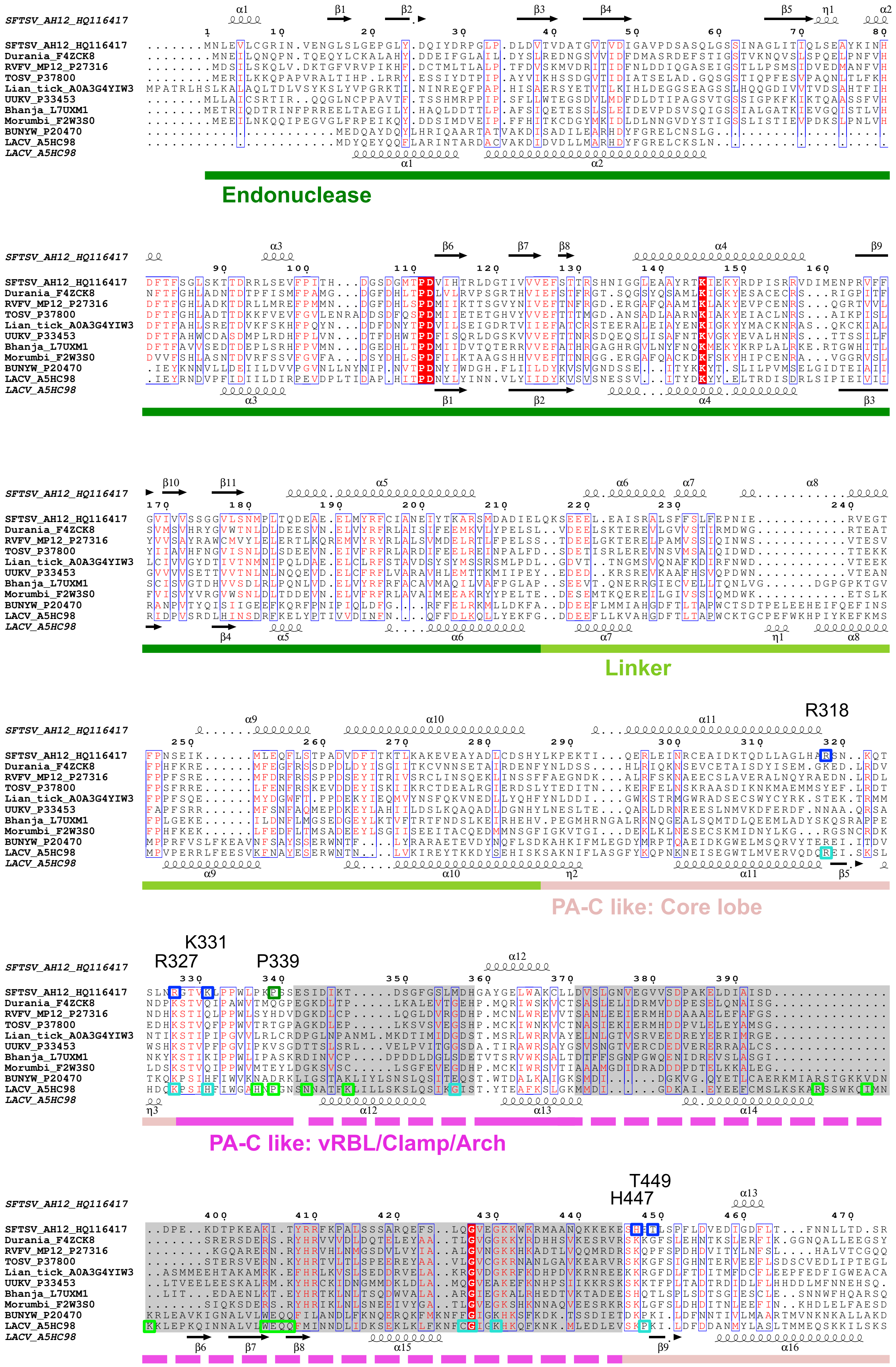
SFTSV_AH12_HQ116417
Durania_F4ZCK8
RVFV_MP12_P27316
TOSV_P37800
Lian_tick_A0A3G4YIW3
UUKV_P33453
Bhanja_L7UXM1
Morumbi_F2W3S0
BUNYW_P20470
LACV_A5HC98
Endonuclease
Linker
R318
PA-C like: Core lobe
K331
R327
P339
PA-C like: vRBL/Clamp/Arch
T449
H447

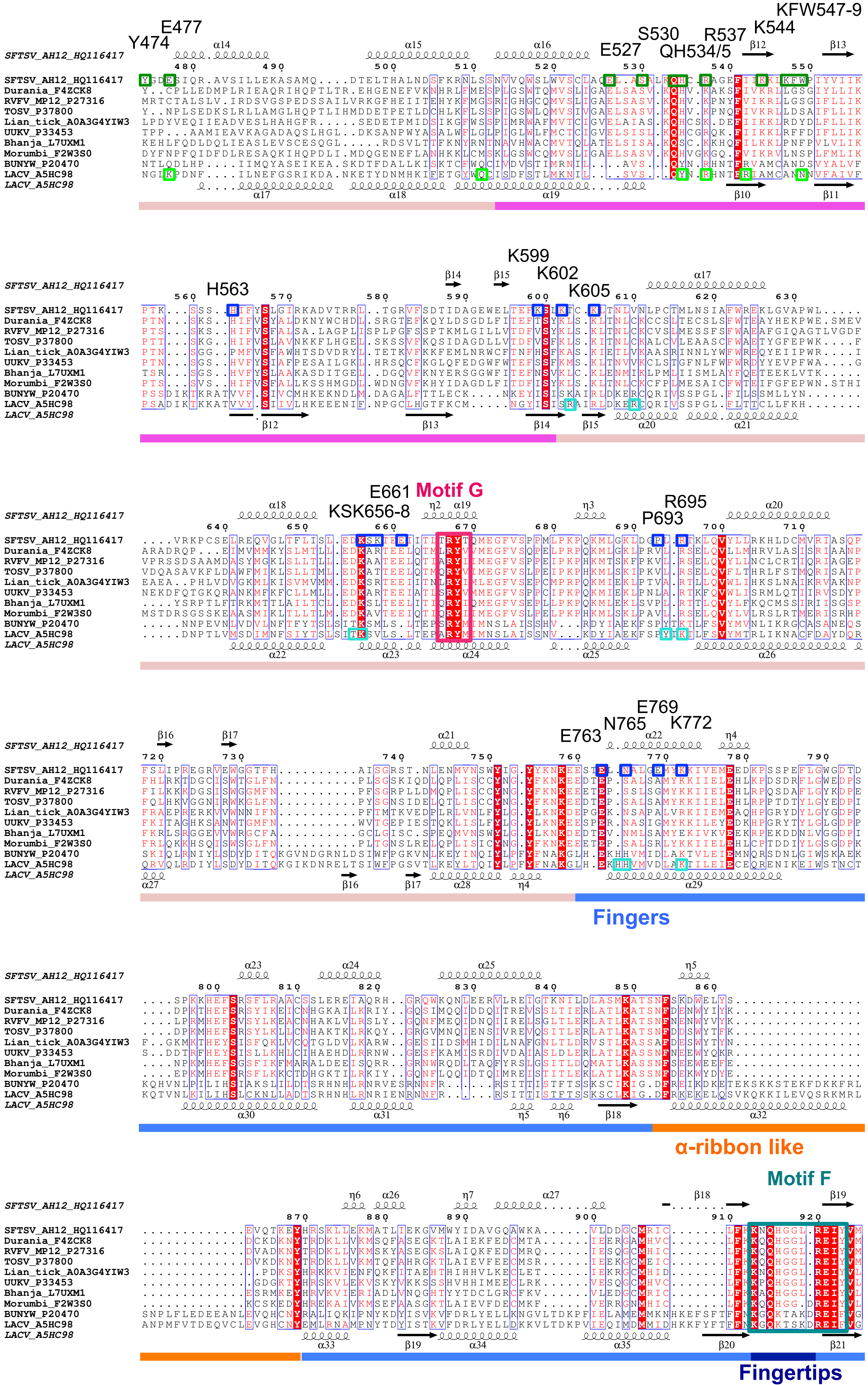

SFTSV_AH12_HQ116417
Durania_F4ZCK8
RVFV_MP12_P27316
TOSV_P37800
Lian_tick_A0A3G4YIW3
UUKV_P33453
Bhanja_L7UXM1
Morumbi_F2W3S0
BUNYW_P20470
LACV_A5HC98
Y474
E477
E527
S530
QH534/5
R537
K544
KFW547-9
H563
K599
K602
K605
E661
KSK656-8
Motif G
R695
P693
E769
N765
K772
E763
Fingers
α-ribbon like
Motif F
Fingertips

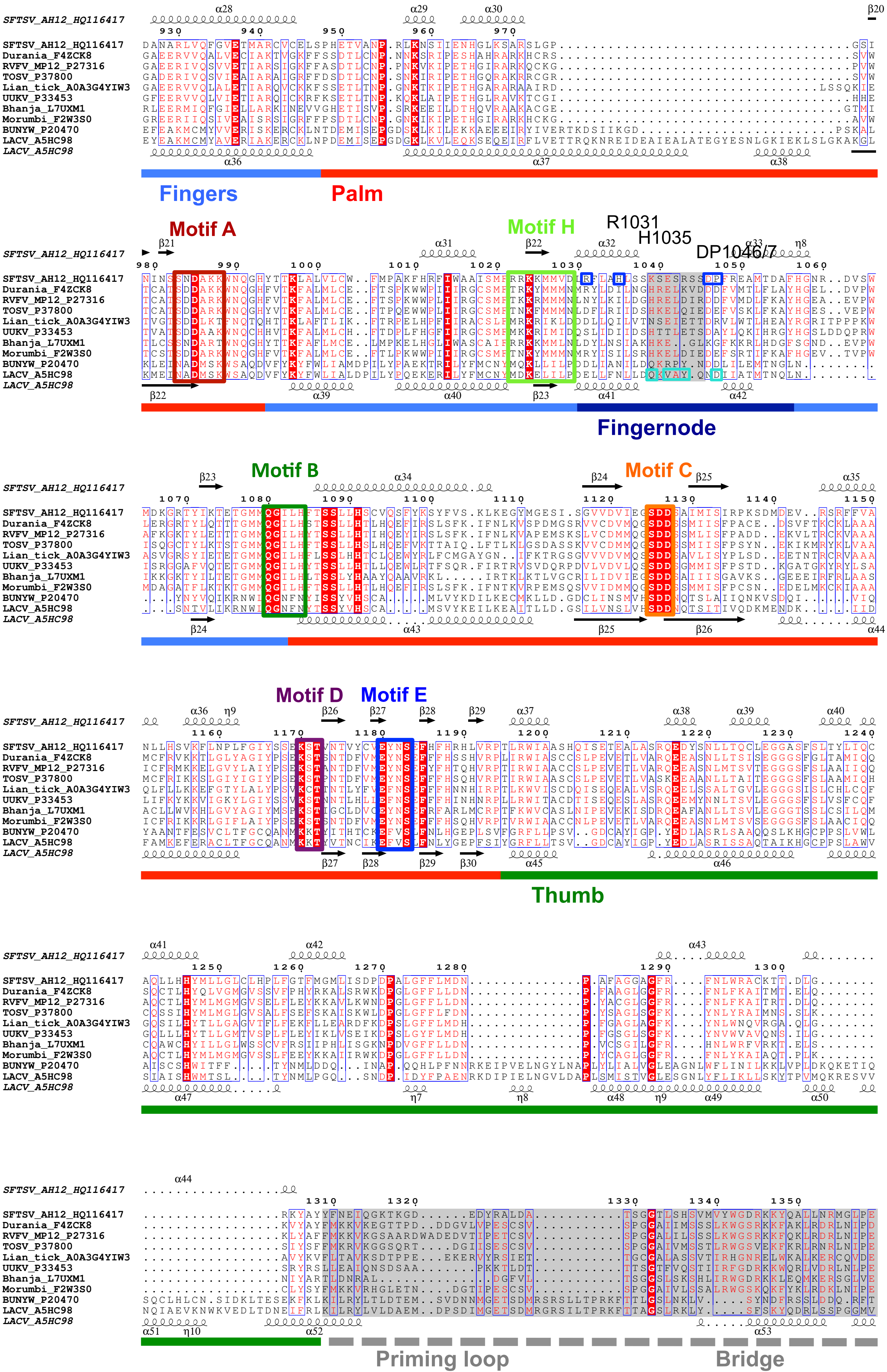

SFTSV_AH12_HQ116417
Durania_F4ZCK8
RVFV_MP12_P27316
TOSV_P37800
Lian_tick_A0A3G4YIW3
UUKV_P33453
Bhanja_L7UXM1
Morumbi_F2W3S0
BUNYW_P20470
LACV_A5HC98
Fingers
Palm
Motif A
Motif H
R1031
H1035
DP1046/7
Fingernode
Motif B
Motif C
Motif D
Motif E
Thumb
Priming loop
Bridge

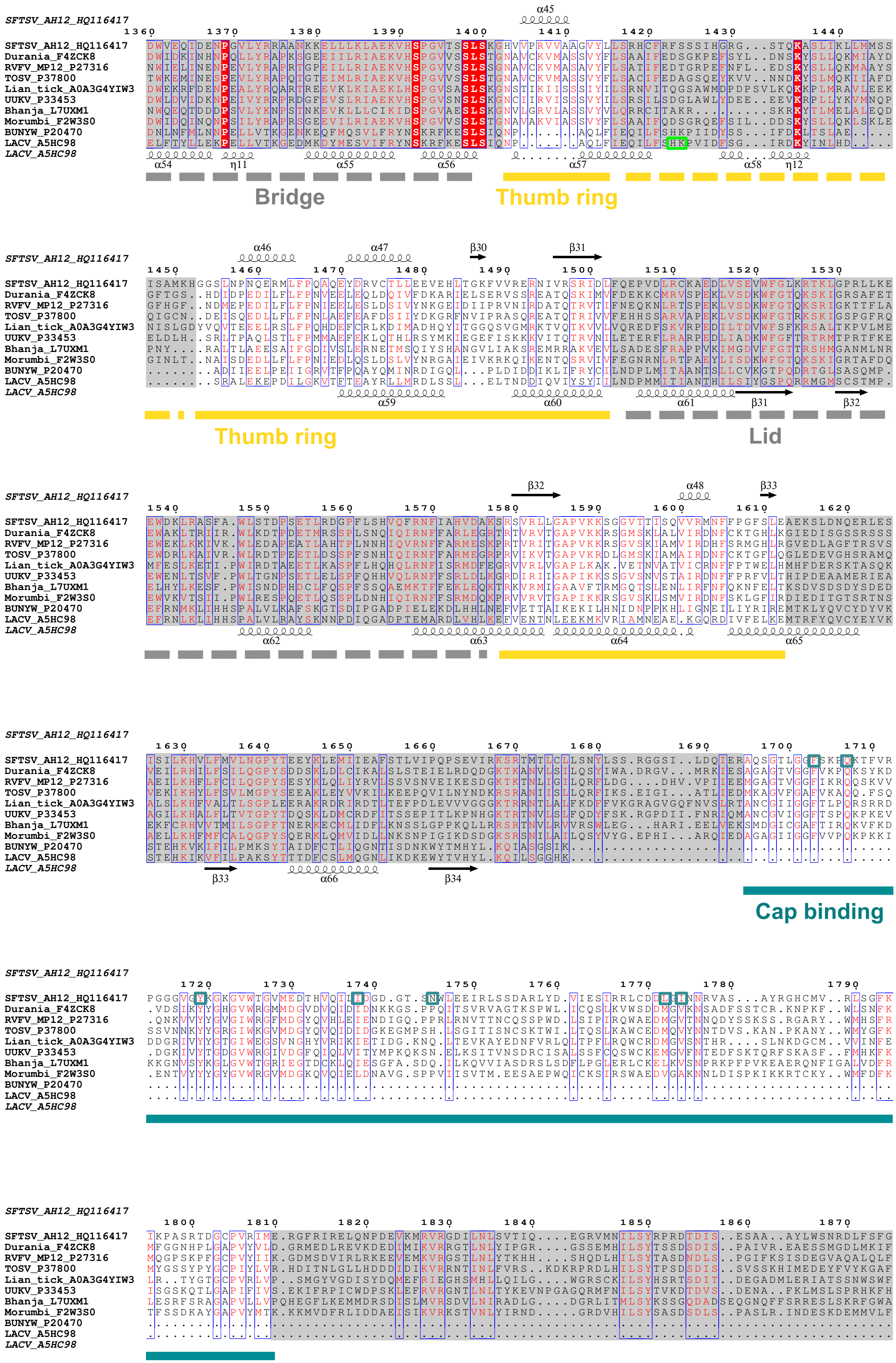
SFTSV_AH12_HQ116417
Durania_F4ZCK8
RVFV_MP12_P27316
TOSV_P37800
Lian_tick_A0A3G4YIW3
UUKV_P33453
Bhanja_L7UXM1
Morumbi_F2W3S0
BUNYW_P20470
LACV_A5HC98
Bridge
Thumb ring
Lid
Cap binding
1360
1370
1380
1390
1400
1410
1420
1430
1440
1450
1460
1470
1480
1490
1500
1510
1520
1530
1540
1550
1560
1570
1580
1590
1600
1610
1620
1630
1640
1650
1660
1670
1680
1690
1700
1710
1720
1730
1740
1750
1760
1770
1780
1790
1800
1810
1820
1830
1840
1850
1860
1870

*SFTSV_AH12_HQ116417*

```
                       1880      1890      1900      1910      1920      1930          1940      1950      1960
SFTSV_AH12_HQ116417    KKEPSCSWICLKTLDNWAWSHASVLLANDRKT.QGIDNRAMGNIFRDCLEGSLRKQG.LMRSKL..TEMVEKNVVPLT.TQELVDILEED
Durania_F4ZCK8         EKEPSKSWICCRSLPVQFVEVLLDVCDGIRKV.DRINPIRLAEIIKICTDSSLRTKVGTVFSMQ..QFTSEAHVVDC...DDLFDVMIED
RVFV_MP12_P27316       EREPSNCWVRCESVAPKFISAILEICEGKRQI.KGINRTRLSEIVRICSESSLRSKVGSMFSFV..ANVEEAHDVDY...DALMDLMIED
TOSV_P37800            SVEPTRSWMLCQPLPWSFVRPVLQVATGSRRSPRQLDLERLREIIRLCTESSIRNKVGTVYGQNRPEKFIEAEPIDM...SEMFDMMLDE
Lian_tick_A0A3G4YIW3   KQEPSHSWYTLTSLNYGTYKFHKEIIEKGGRL.GNIDIQAYREILKQSAHYTLAMRGFTMHDIP.PQTITSGRARDF...DKQMWKMIKE
UUKV_P33453            GREPSTSWICMRALPISTIDKLLERILNRERISGSIDNERLAECFKNVMESTLRRKGVFLSEFS..RATQKMLDGLS...RDMLDFFAEA
Bhanja_L7UXM1          IREPYLSWISLGPIPDVLVPRLKEEMTGRRRT.EGIDGEQLRDIFKRCCKGSLRRKGLTVGQYS..NVRLRIEDTSEEVDIDLLELLDED
Morumbi_F2W3S0         NKEPSKSWASCSPIPAIMMHKIIKVVQGELKI.ASLDSSRLGEIMKLCCESSLRSRIGTLFSAL..PSVQNTSRVDV...DDLIDIVLTD
BUNYW_P20470           ..........................................................................................
LACV_A5HC98            ..........................................................................................
```

*LACV_A5HC98*

*SFTSV_AH12_HQ116417*

```
                                   1970      1980      1990      2000      2010      2020      2030
SFTSV_AH12_HQ116417    IDF......SDVIAVELSEGSL..DIES.IFD.GAPILWSA..EVEEFGEGVVAVSYSSKYYHLT.LMDQAAITMCAIMGKEGCRGLLTE
Durania_F4ZCK8         MNTGIFGEVVNKMEEEVRGDNE...DLE..FDFEDINLFGPA.HFKES........SELSLVSHP.LMDRFVDSAIAKMGRSAMRRLLEV
RVFV_MP12_P27316       AKNNAFSHVVDCIELDVSGPYE...MES..FHGRSTLTCTPSTILIRT........YTFYLTLHQ.TMIS.VQAFQVILDEGVLLIALVN
TOSV_P37800            GMDDAFEELADYLTVEEDPDYM..DEVS..FDDDSLNLFGPA.HYKEL........QSLTVLAHP.LMDDFVTRLVGKMGRPQIRRLLEK
Lian_tick_A0A3G4YIW3   GAM...SKMFETE.....QARA..LLEGAEFEEGEDRQEAGPSTPRKDEDSRRIAIGLFPMGVNKGRIMESIDVDLFGEP.DQPIIPATL
UUKV_P33453            GLN...DDLLLEEEPWLSGLDTFMLDDEAYLEEYNLGPFGVF.SVEQE........MNTKYYHHL.LLDSLVEDVIQKLSLDGLRKLFQE
Bhanja_L7UXM1          VDF...GDTLQSQNIQDLEASI..EEMEVLFGEDDLEVFDFT.EVISR........SDPL.AYHG.FCDRIIEDYIKRLGHEAIRKAIQK
Morumbi_F2W3S0         SKTAGFREIVQSLENDIKDEYE...VED..FDLSDIDLFGPA.HYKEL........SDLNTISHP.LMDDYVEFCISTIGRKELRRVLET
BUNYW_P20470           ..........................................................................................
LACV_A5HC98            ..........................................................................................
```

*LACV_A5HC98*

*SFTSV_AH12_HQ116417*

```
                      2040      2050      2060          2070          2080
SFTSV_AH12_HQ116417    KRCMAAIREQVRPFLIFLQIPEDSIS.WVS.DQFC...DSRGLDEESTIMWG
Durania_F4ZCK8         GYTPTRYIHVSRLIYRAIGRNPDSIK.TDD.LEIL..QDMDISDE.MIG...
RVFV_MP12_P27316       NYLRGSKANCWVRCESVAPKFISAILEICEGKRQIKGINRTRLSEIVEFVLN
TOSV_P37800            NVTTRDLRELSELLFMALDRDPSQIR.EEL.ILGD...SPTEVPDD.LLG..
Lian_tick_A0A3G4YIW3   FRHPTLSANCVDNIIG............TEMT.....PTEVTAVLEKW....
UUKV_P33453            EEAPLEYKKEVIRLLNILQRDASQIK.WKS.RDLLSENMGLDVDDD......
Bhanja_L7UXM1          RECAQSDYPLVKNLFESIGENPDELSVLPDKDDAY.ASSG.LITEDLWG...
Morumbi_F2W3S0         NRCKTKDLQLSKDLFLVLGRNPDDIK.VDE.YNLR...EQMAVEDDMIG...
BUNYW_P20470           ....................................................
LACV_A5HC98            ....................................................
```

*LACV_A5HC98*
